# Atlas-Guided Discovery of Transcription Factors for T Cell Programming

**DOI:** 10.1101/2023.01.03.522354

**Authors:** H. Kay Chung, Cong Liu, Anamika Battu, Alexander N. Jambor, Brandon M. Pratt, Fucong Xie, Brian P. Riesenberg, Eduardo Casillas, Ming Sun, Elisa Landoni, Yanpei Li, Qidang Ye, Daniel Joo, Jarred Green, Zaid Syed, Nolan J. Brown, Mattew Smith, Shixin Ma, Shirong Tan, Brent Chick, Victoria Tripple, Z. Audrey Wang, Jun Wang, Bryan Mcdonald, Peixiang He, Qiyuan Yang, Timothy Chen, Siva Karthik Varanasi, Michael LaPorte, Thomas H. Mann, Dan Chen, Filipe Hoffmann, Josephine Ho, Jennifer Modliszewski, April Williams, Yusha Liu, Zhen Wang, Jieyuan Liu, Yiming Gao, Zhiting Hu, Ukrae H. Cho, Longwei Liu, Yingxiao Wang, Diana C. Hargreaves, Gianpietro Dotti, Barbara Savoldo, Jessica E. Thaxton, J. Justin Milner, Susan M. Kaech, Wei Wang

## Abstract

CD8^+^ T cells differentiate into diverse states that shape immune outcomes in cancer and chronic infection^1–4^. To systematically define the transcription factors (TFs) driving these states, we built a comprehensive atlas integrating transcriptional and epigenetic data across nine CD8^+^ T cell states and inferred TF activity profiles. Our analysis catalogued TF activity fingerprints, uncovering regulatory mechanisms governing selective cell state differentiation. Leveraging this platform, we focused on two transcriptionally similar but functionally opposing states critical in tumor and viral contexts: terminally exhausted T cells (TEX_term_), which are dysfunctional^5–8^, and tissue-resident memory T cells (T_RM_), which are protective^9–13^. Global TF community analysis revealed distinct biological pathways and TF-driven networks underlying protective versus dysfunctional states. Through *in vivo* CRISPR screening integrated with single-cell RNA sequencing (*in vivo* Perturb-seq), we delineated several TFs that selectively govern TEX_term_ differentiation. We also identified HIC1 and GFI1 as shared regulators of TEX_term_ and T_RM_ differentiation and KLF6 as a unique regulator of T_RM_. Importantly, we discovered novel TEX_term_- selective TFs, including ZSCAN20 and JDP2, with no prior known function in T cells. Targeted deletion of these TFs enhanced tumor control and synergized with immune checkpoint blockade but did not interfere with T_RM_ formation. Consistently, their depletion in human T cells reduces the expression of inhibitory receptors and improves effector function. By decoupling exhaustion T_EX_-selective from protective T_RM_ programs, our platform enables more precise engineering of T cell states, accelerating the rational design of more effective cellular immunotherapies.

---

Cell states are the range of cellular phenotypes arising from a defined cell type’s interaction with its environment. Within the immune system, T cells possess multiple differentiation states, particularly as naive T cells differentiate into diverse states with different functionality and trafficking patterns in various immune environments, such as tumors and virus infections^1–4^. As transcription factors (TFs) govern cell state differentiation^14^, understanding how TFs shape these states is essential for programming beneficial states with therapeutic potential. One promising application of cell state engineering is enhancing CD8^+^ T cells for adoptive cell transfer therapy (ACT) of tumor-infiltrating lymphocytes (TILs) or CAR-T cells. However, identifying TFs that control CD8⁺ T cell states is difficult due to substantial heterogeneity and overlapping transcriptomes, even between functionally divergent states.

We focused on two transcriptionally similar yet functionally divergent states: the protective tissue-resident memory (T_RM_) cell state and the dysfunctional terminally exhausted (TEX_term_) cell state. Many studies show that TILs with T_RM_ characteristics correlate with better survival in solid tumor patients^9–13^. Conversely, during persistent antigen stimulation scenarios such as chronic virus infection (e.g., HIV) or cancer, T cells progressively express diverse inhibitory receptors, including PD1, lose memory potential, and effector functions. This process is known as T cell exhaustion (TEX)^5,7,8,15^, and cells in this trajectory eventually adopt the TEX_term_ cell state. TEX_term_ cells express higher levels of diverse inhibitory receptors (e.g., TIM3 and CD101), lack effector and proliferative capacity, and do not respond effectively to immune checkpoint blockade (ICB), such as anti-PD1 monoclonal antibody (mAb) blockade^16–18^. High TEX_term_ marker expression often indicates poor prognosis in solid tumors, though some markers also correlate with ICB response, highlighting their complex role in tumor immunity^19,20^. Despite their distinct functional impact on cancer outcomes, TEX_term_ and T_RM_ cells both preferentially reside in tissues^1,3^ and display remarkable similarities in their transcriptional profiles, including key regulatory TFs such as BLIMP1^5,21–23^, BHLHE40^24,25^, and NR4A2^9,26,27^(Fig. 1a,b, Extended Data Fig. 1a-c). These two cell states even exhibit highly correlated open chromatin regions (Extended Data Fig. 1d), complicating the precise identification of TFs whose disruption may selectively inhibit TEX_term_ while preserving T_RM_ cell development. Given that many TFs are commonly expressed across different CD8^+^ T cell states and differentiation trajectories, a sophisticated and precise bioinformatics approach is crucial to pinpoint the bona fide cell-state-specifying TFs essential for T cell programming.

**Fig. 1.**
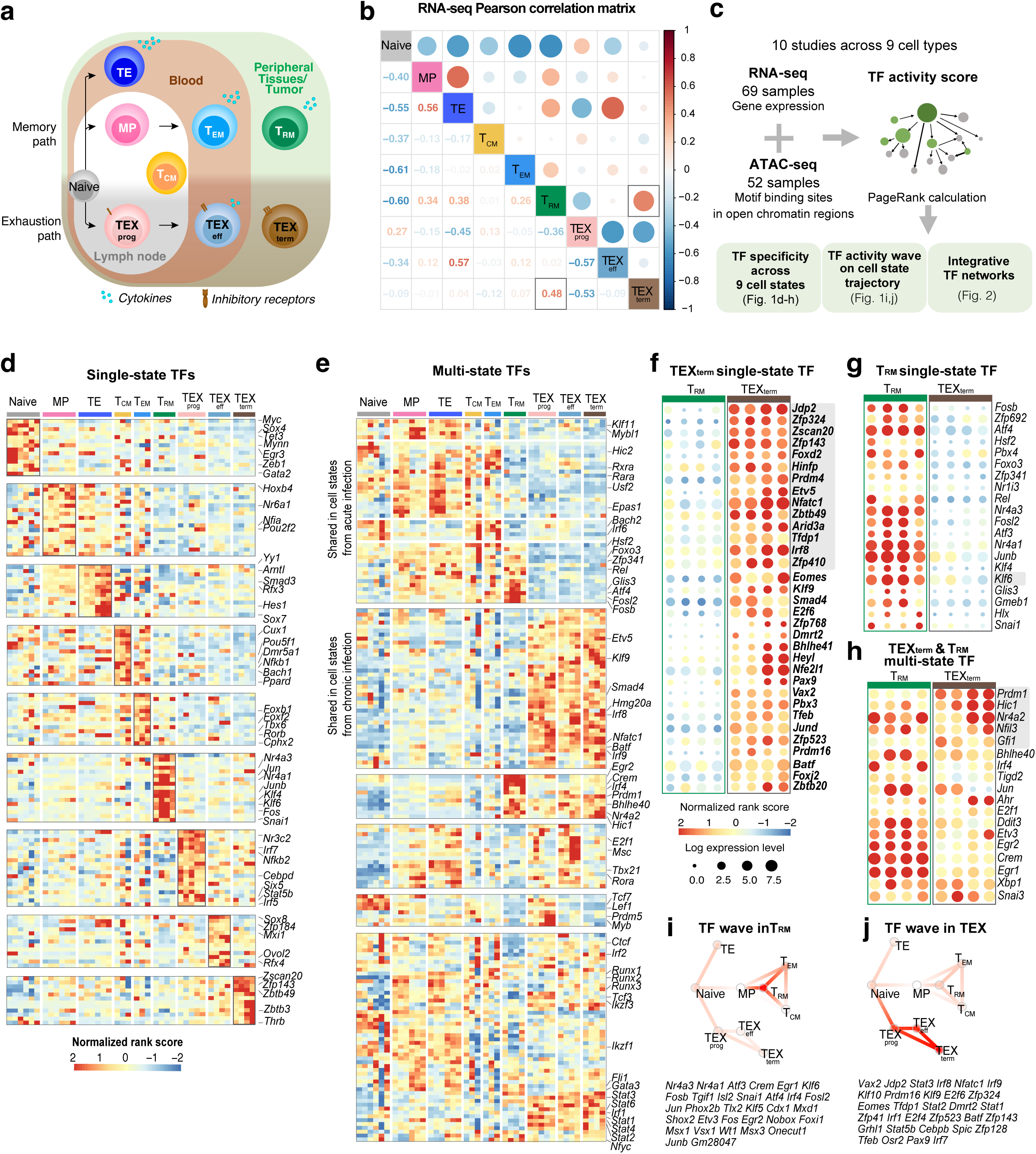
| **Transcriptional and epigenomic atlas of CD8^+^ T cell differentiation states and TF identification pipeline. a,** Diagram summarizing CD8⁺ T cell trajectories during acute and chronic infection or tumor, highlighting differentiation into various effector, memory, and exhaustion states, including parallel TRM and TEXterm lineages with overlapping tissue localization. **b,** Pearson correlation matrix of batch-effect–corrected RNA-seq datasets. Both color intensity and circle size indicate correlation strength, with red denoting the highest correlation. **c,** Workflow of the integrative Taiji analysis. Matched RNA-seq and ATAC-seq datasets^3,9,18,23,32–36^ were used to construct a regulatory network and calculate TF activity scores using PageRank. Downstream analysis included identification of single- and multi-state TFs, TF “waves” and network communities. **d-h,** TFs (rows) and samples (columns) are displayed as z-normalized PageRank heatmaps. Each column corresponds to a dataset. **d,** PageRank scores of 136 single-state TFs. **e,** 173 multi-state TFs. **f-h**, Bubble plots show normalized TF PageRank scores and expression for TEX_term_-selective **(f),** T_RM_ -selective **(g),** and multi-state **(h)** TFs active in both cell states. TEX_term_ single-state TFs are bolded. Circle color represents the normalized PageRank score (red = high), and circle size indicates log mRNA expression across five datasets. TFs are ordered by *P* value; validated TFs are highlighted in gray. **i-j,** TF “waves” associated with exhaustion **(i)** or T_RM_ differentiation **(j)**, indicating coordinated activity of TF groups during cell state transistions. Sample sizes and statistical details for cell state definitions and TF selection criteria are provided in Extended Data Fig. 1e and Extended Data Fig. 2a, respectively.

We hypothesized that identifying key TFs controlling selective CD8⁺ T cell differentiation can be achieved through systematic comparison of TF activity across the differentiation landscape. Accurate prediction requires recognizing that TF activity does not necessarily mirror expression, as it depends on post-translational modifications, cofactors, and target accessibility^28^, and that TF effects propagate through genetic networks. We therefore developed a multi-omics atlas integrating transcriptomic and chromatin data from nine CD8⁺ T cell states to understand “global” influences of TFs in each cell state and to identify “*selective*” or “*shared*” TFs. Our atlas-based platform can map TF communities and their target genes (“regulatees”), guiding state-specific differentiation.

## Multiomics atlas maps of CD8⁺ T Cell TFs

Our initial objective was to create a comprehensive catalogue of TF activity across diverse CD8^+^ T cell states by integrating our TF activity analysis pipeline, Taiji^29–31^, with comparative statistical analysis. In Taiji, the gene regulatory network (GRN) is a weighted, directed network that models regulatory interactions between TFs and their target genes. In this GRN, each node corresponds to a gene, and its weight is proportional to the gene’s expression level. Each edge represents a regulatory interaction and is weighted based on a combination of factors: the TF’s predicted binding affinity to the target gene, the chromatin accessibility at the target gene’s locus, and the expression levels of both the TF and the target gene (Fig. 1c)^29,30^. To determine the global influence of each TF within the network, Taiji applies a personalized PageRank algorithm, which assigns an “importance” score to each node based on both the quantity and quality of incoming connections. This approach yields a measure of TF activity that reflects the influence of each TF in the broader regulatory landscape, accounting for upstream regulators, downstream targets, and feedback loops through iterative computation.

With Taiji, we previously identified TFs involved in pan-immune lineage commitment, including NK cells, dendritic cells, B cells, and γδ T cells^35^. While earlier studies provided foundational insights into cell differentiation, a more refined analysis within CD8^+^ T cells is needed to achieve higher resolution of TF roles. Therefore, leveraging the improved statistical filtering, we aimed to quantify the global influence of TFs across all CD8^+^ T cell states.

To begin, we analyzed ATAC-seq and RNA-seq datasets from 121 CD8^+^ T cell samples spanning nine distinct states, using both previously published and newly generated datasets from well-characterized acute and chronic lymphocytic choriomeningitis virus (LCMV) infections^3,9,18,23,32–36^ (Extended Data Fig. 2; Supplementary Table 1). In acute LCMV-Armstrong infections, CD8^+^ T cells differentiate into memory precursor (MP), terminal effector (TE), effector memory (T_EM_), central memory (T_CM_), and T_RM_ states. In chronic LCMV-Clone 13 infections, they adopt heterogeneous exhaustion cell states, including progenitors of exhaustion (TEX_prog_), effector-like exhaustion (TEX_eff_), and TEX_term_.

Next, we conducted an unbiased comparative analysis using statistical filtering to understand the specificity of TF activity across the CD8^+^ T cell states (Extended Data Fig. 2a, Supplementary Table 2). This identified TFs; 136 were predominantly ‘**single-state**’ TFs, with each cell state selectively containing 12-19 unique TFs (Fig. 1d, Extended Data Fig. 2b). This category included novel TFs such as *Hoxa7* in Naïve, *Snai1* in T_RM_, *Hey1* in TEX_prog_, *Sox8* in TEX_eff_, and *Zscan20* and *Jdp2* in TEX_term_ cells. In contrast, 119 TFs were key regulators in more than one cell state, termed ‘**multi-state’** TFs (Fig. 1e), such as *Tcf7* and *Tbx21*. *Tcf7* is a known driver of Naive-, MP-, and TEX_prog_ states, all multipotent with high proliferative capacity^3,18^. Multi-state TFs like *Vax2*, *Batf*, *Irf8,* and *Stat1* were more enriched within the exhaustion-associated cell states (TEX_prog_, TEX_eff_, and TEX_term_). Consistent with TEX_term_ and T_RM_ similarity (Fig. 1b; Extended Data Fig. 1b-d), both cell states share the most TFs compared to other cell states (e.g., *Egr2*, *Crem*, *Prdm1*, Extended Data Fig. 2c).

While Taiji provides a statistically grounded approach for inferring TF activity, there is no absolute threshold for defining cell state specificity, and some misclassification is expected, particularly for TFs with overlapping functions or modest differences in activity. Still, Taiji is useful to highlight TFs with activity patterns enriched in specific cell states. For instance, although *Eomes* was classified as a TEX_term_ single-state TF herein, it also functions in effector, T_EM_, T_CM_, and T_RM_ differentiation^37,38^. This illustrates that more accurate classifications require further investigation and resolution, as performed herein for several TFs.

## TF state-selectivity in TEX_term_ and T_RM_ cells

Despite the strong transcriptional overlap between TEX_term_ and T_RM_ cells, our Taiji pipeline predicted TFs selectively active in either of these two cell states. This could aid in developing better immunotherapies, in which one can engineer T cells away from exhaustion and toward more functional effector cell states without negatively impacting T_RM_ formation in tissues and tumors. Based on statistical criteria (Extended Data Fig. 2a), we identified 20 and 34 TFs as single-state TFs of T_RM_ or TEX_term_ cells, respectively, and 30 multi-state TFs active in both (Fig. 1f–h; Extended Data Fig. 2a, blue boxes; Supplementary Table 3). TEX_term_ single-state TFs included many previously unreported TFs such as *Zscan20*, *Jdp2*, *Zfp324*, *Zfp143*, *Zbtb49*, and *Arid3a* (Fig. 1f). T_RM_ single-state TFs included *Fosb*, *Zfp692*, *Atf4*, *Pbx4, Junb, and Klf6* (Fig. 1g). Of the TEX_term_ and T_RM_ multi-state TFs, some were well-known to function in the development of *both* cell states, such as *Nr4a2*^12^, *Bhlhe40*^24^, *Prdm1*^23,35^, whereas others such as *Hic1*^39^ and *Gfi1*^40^ were not, identifying them as new multi-state TFs to consider (Fig. 1h). We analyzed previously reported TFs such as cJUN, BATF/BATF3, and TFAP4 that were identified from functional screening of CD8^+^ T cells^41–44^ based on limited phenotypic readouts. These prior screens tended to identify broadly active, multi-state TFs (see Fig. 1). In contrast, our platform enabled a computationally guided, multi-state screen that revealed TFs predicted to have greater state-selective activity (Extended Data Fig. 2d).

To evaluate the predicted TFs governing selective T cell differentiation, we identified dynamic activity patterns of TF groups, termed "TF waves” (Extended Data Fig. 3). TF waves reveal possible combinations of TFs that coordinate trajectories. Seven TF waves linked to specific biological pathways were identified, such as the T_RM_ TF wave (Fig. 1i), including several members of the AP-1 family (e.g., *Atf3*, *Fosb*, and *Jun*) is uniquely associated with the TGFβ response pathway (Extended Data Fig. 4e). The TEX TF wave, involving *Irf8*, *Jdp2*, *Nfatc1*, and *Vax2*, correlates with PD1 and senescence pathways (Fig. 1j, Extended Data Fig. 3e).

## TF community analysis of T_RM_ vs TEX_term_ cells

To uncover transcriptional programs governing T_RM_ or TEX_term_ differentiation, we constructed TF–TF association networks capturing functional relationships between TFs (Fig. 2a). Analysis of regulatee- based adjacency matrices (i.e., predicted TF-target gene circuits) revealed shared and distinct patterns of TF collaboration across the two states. Single-state TFs displayed strong intra-state connectivity. TEX_term_ TFs (ZSCAN20, JDP2, ZFP324, IRF8) formed dense networks within TEX_term_ cells (Fig. 2b; Extended Data Fig. 4a), while T_RM_ TFs (FOSB, SNAI1, KLF6) interacted mainly within T_RM_ networks Extended Data Fig. 4b). Interestingly, multi-state TFs (HIC1, PRDM1, FLI1, GFI1), active in both states and previously reported TFs (cJUN, BATF, and TFAP4) formed distinct partnerships in each cell state, reflecting context-specific regulatory architectures (Fig. 2c; Extended Data Fig. 4c).

**Fig. 2.**
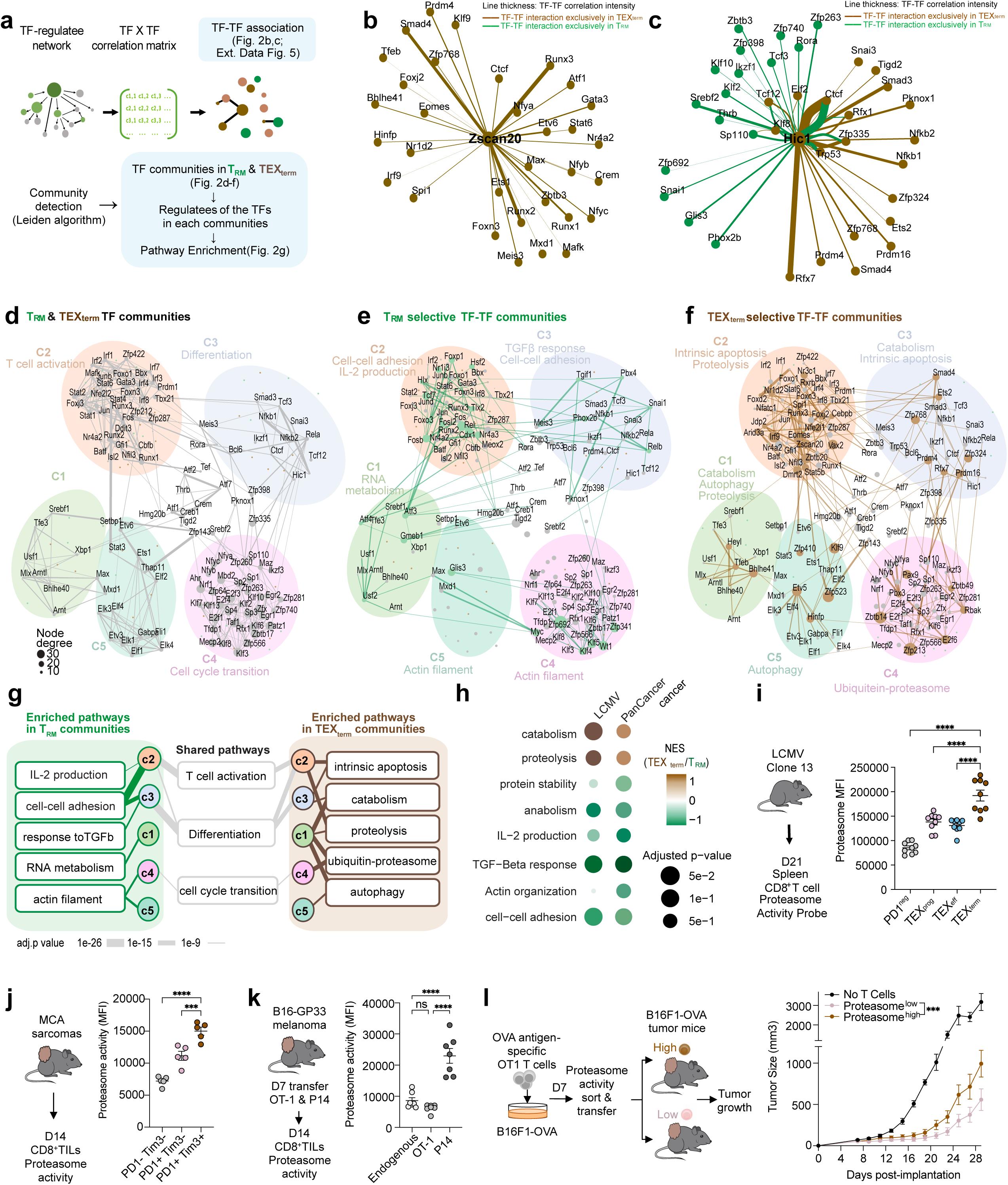
| **Global analysis of TF networks in TEX_term_ and T_RM_ cell states. a,** Overview of TF–TF network analysis encompassing association and community-level organization of TRM and TEXterm regulatory landscapes. **b-c,** TF-TF association networks focused on the TEX_term_ single-state TF ZSCAN20 **(b)** and the multi-state TF HIC1 **(c)**, depicting predicted context-specific interactions in T_RM_ (green) or TEX_term_ (brown) cells. **d-f,** Clustering of TF–TF associations identified five distinct TF communities in TRM and TEXterm networks. Shared TFs (gray) shape overall community topology **(d),** whereas T_RM_- or TEX_term_-specific interactions are represented as green **(e)** or brown **(f)** edges, respectively. **g,** Summary of shared and unique biological pathways enriched within T_RM_ and TEX_term_ communities. Line thickness reflects −𝑙𝑜𝑔_10_(*P* value). (Pathway gene sets in Supplementary Table 8). **h**, Gene set enrichment analysis (GSEA) comparing TEX_term_ vs. T_RM_ cell pathways using batch-effect corrected LCMV bulk- RNA-seq^3,9,18,23,32–36^ and human pan-cancer scRNA-seq data sets^45,56,61^. **i-k,** Flow cytometry analysis of proteasome activity showing the highest activity in TEX_term_ cells during LCMV-Clone 13 infection **(i)** and MCA-205 tumors **(j).** In dual transfer experiments, antigen-specific (P14) and bystander (OT-1) CD8^+^ T cells analyzed from B16-GP33 tumors **(k)** show elevate proteasome activity in TEX_term_-like populations. **l,** Functional impact of proteasome activity on tumor growth. Tumor-bearing C57BL/6 mice infused with proteasome^high^ or proteasome^low^ OT-1 cells pre-stimulated with B16F1-OVA tumor cells for seven days. Proteasome^high^ OT-1 cells exhibit reduced tumor control. Data are shown as the mean ±s.e.m. Ordinary one-way ANOVA **(i-k),** and two-way ANOVA Tukey’s multiple comparison test **(l)** were performed. *\*\*\*\*P*<0.0001, *\*\*\*P*<0.001, *\*\*P*<0.01, *\*P*<0.05.

We next grouped the TF–TF association networks into distinct “TF neighbor communities” in T_RM_ and TEX_term_ cells (Supplementary Table 5), and each community was linked to specific biological processes (Fig. 2d–f). While multi-state TFs shaped overall community topology, single-state TFs drove unique interaction patterns specific to T_RM_ or TEX_term_ cells within each community. Pathway analysis revealed divergent programs in each state—for instance, T_RM_ community-3 was associated with cell adhesion and TGFβ response (Fig. 2e, g, h), while TEX_term_ community-3 was linked to apoptosis (Fig. 2f-h). Community-1 in T_RM_ cells controlled RNA metabolism (Fig. 2e, g, h), whereas in TEX_term_ cells, it was tied to catabolism, proteolysis, and autophagy (Fig. 2f-h).

To assess the functional relevance of state-enriched pathways, we focused on the proteasome pathway, which emerged as a prominent but previously unrecognized feature of TEX_term_ cells (Fig. 2g, h). Proteasome gene signatures were enriched in TEX_term_-like CD8⁺ T cells from NSCLC patients^45^ and murine MCA-205 TILs (Extended Data Fig. 5a,b). Consistently, proteasome activity—measured by a validated fluorescent probe^46^—was highest in TEX_term_ cells from chronic LCMV (Fig. 2i) and in tumor- specific TILs (Fig. 2j, k) relative to bystander OT-1 cells (Fig. 2k). To test if high proteasome activity correlates with dysfunction, we sorted OT-1 cells by proteasome activity probe intensity and adoptively transferred them into B16F10-OVA tumor-bearing mice. Proteasome^high^ cells showed reduced tumor control compared to proteasome^low^ cells (Fig. 2l), a trend also seen in endogenous TILs (Extended Data Fig. 5c). These findings support the TF-TF network and pathway predictions and identify the proteasome pathway as a functional hallmark of TEX_term_ cells.

### *In vivo* CRISPR Screens of TEX_term_ TFs

The Taiji pipeline enabled comparative analysis of TF activity and curated sets of single-state TFs specific to T_RM_ *vs.* TEX_term_ cells (Fig. 1f-h). To assess its accuracy, Perturb-seq, combining *in vivo* CRISPR screening with single-cell RNA sequencing (scRNA-seq), was performed in two different animal models for T_RM_ or TEX_term_ differentiation (Fig. 3a and 4a). Our Perturb-seq guide RNA (gRNA) library targeted 19 TFs, including seven TEX_term_ and T_RM_ multi-state TFs and 12 TEX_term_ single-state TFs. The TEX_term_-TFs included one known TF, *Nfatc1*, and 11 others with high specificity scores but not previously linked to TEX_term_ differentiation (gray boxes Fig. 1f,h). The multi-state TFs included two positive controls (*Nr4a2*, *Prdm1*) and unvalidated multi-state TFs (*Nfil3*, *Hic1*, *Gfi1*, *Ikzf3*, *Stat3*. To ensure comprehensive screening, four gRNAs per target were expressed in two dual-gRNA retroviral vectors (Extended Data Fig. 6a), along with two control vectors with scramble gRNAs (gScramble). This created a library of 40 dual-gRNA vectors, with 76 TF-gRNAs and 4 gScramble controls (Supplementary Table 6).

**Fig. 3.**
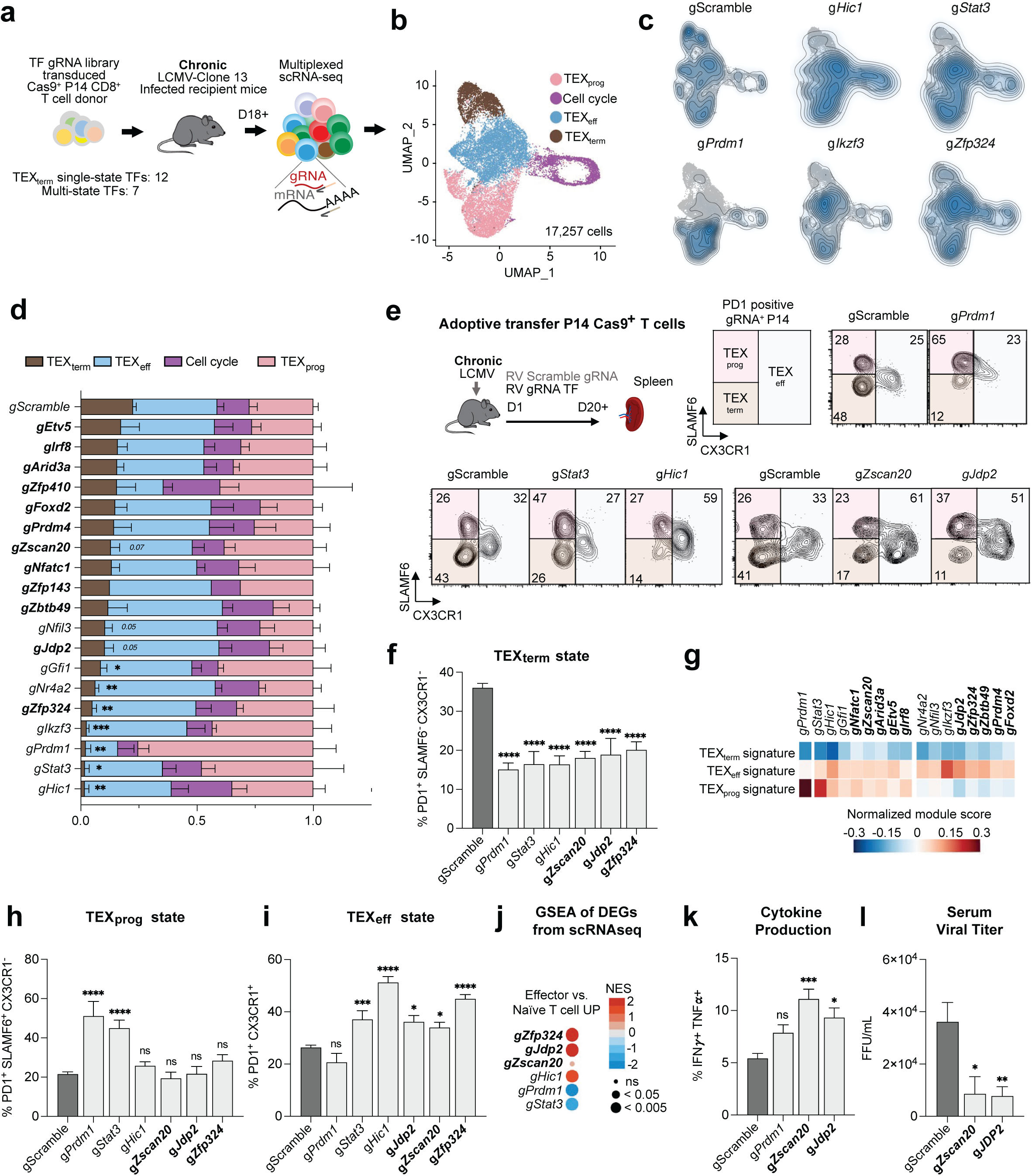
| ***In vivo* Perturb-seq validation of TEX_term_-driving TFs. a**, Schematic of the *in vivo* Perturb- seq strategy. Cas9^+^ P14^+^ TCR-transgenic CD8^+^ T cells recognizing LCMV epitope GP33-41 epitope were transduced with retroviruses (RVs) expressing gRNA libraries, adoptively transferred into mice previously infected one day earlier with LCMV-Clone 13, and analyzed 18-23 days later by scRNA- seq. **b**, UMAP showing TEX_prog_, TEX_eff_, TEX_term_, and cell cycle clusters; marker expression are in Extended Data Fig. 6b,c. **c-d**, Kernel density plots and distributions of gRNA⁺ cells across clusters. TEX_term_ single-state TFs are indicated bolded. Data represent five pooled replicates from three independent experiments; values are shown as mean ± s.e.m. Statistical analysis: two-way ANOVA with Fisher’s LSD test compared to the gScramble control; results for TEX_term_ and TEX_prog_ clusters are shown; with full comparisons in Supplementary Table 7. *\*\*\*\*P* < 0.0001, *\*\*\*P* < 0.001, *\*\*P* < 0.01*, *P* < 0.05. **e**, Representative flow plots showing phenotyping of the TF knockouts (Kos) in LCMV- Clone 13-infected mice. **f**, Quantification of TEX_term_ (PD1^+^ SLAMF6^-^ CX3CR1^-^) frequencies in donor CD8^+^ T cells. **g**, Differential expression analysis of TEX_term_, TEX_prog_, and TEX_eff_ gene signatures (ref. ^47^; Supplementary Table 8) across each TF KO. **h-i**, Frequencies of TEX_prog_ (PD1^+^ SLAMF6^+^ CX3CR1^-^) and TEX_eff_ (PD1^+^ CX3CR1^+^) subsets. **j**, GSEA showing enrichof effector-associated gene sets in TF KOs *vs*. control. **k-l**, Functional validation: cytokine production (IFNγ, TNFα) and viral titers in mice receiving TF KO versus control CD8^+^ T cells. Statistic analysis for **f, h, i, k, l**: mean ± s.e.m., ordinary one-way ANOVA with Dunnett’s multiple comparison versus gScramble (**f-k**: n ≥ 8, ≥ 3 biological replicates, **i**: n ≥ 4, ≥ 2 biological replicates). ***P<0.001, ** P <0.01, * P <0.05.

**Fig. 4.**
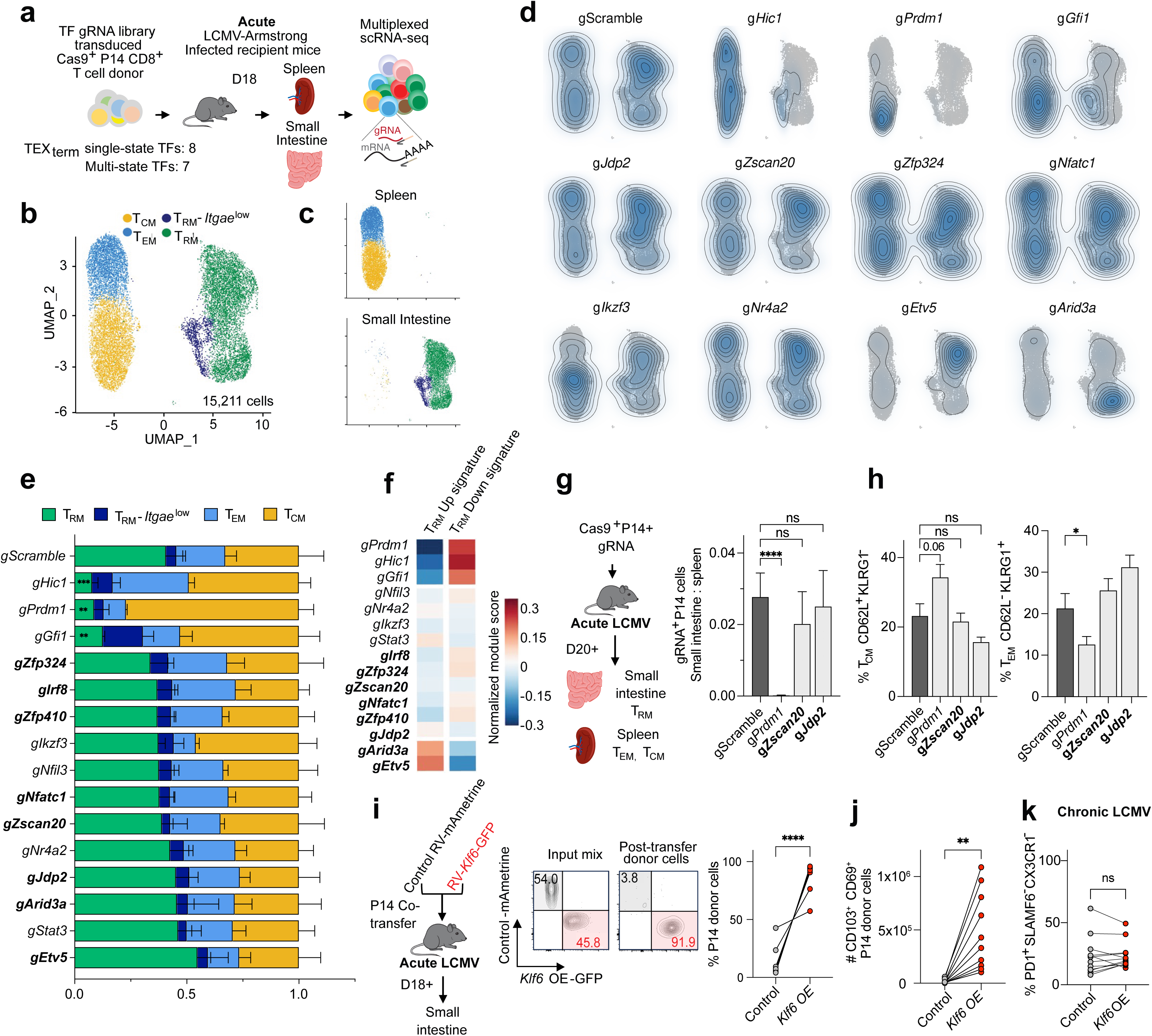
| **Functional validation of TFs with distinct roles in TEX_term_ and TRM differentiation. a,** Schematic of *in vivo* Perturb-seq screening during acute LCMV-Armstrong infection to assess memory CD8^+^ T cell differentiation. Transduced donor Cas9^+^ P14^+^ CD8^+^ T cells were analyzed for tissue-resident (T_RM_), effector-memory (T_EM_), and central-memory (T_CM_) states in the small intestine (SI) and spleen. **b,** UMAP embedding of 15,211 cells identifying T_CM_ (*Il7r*, *Tcf7*, *Sell*, *S1pr1*), T_EM_ (*Cx3cr1*, *Klrg1*, *Klf2*), T_RM_ (*Cd69*, *Itgae*, *Cd160*), and T_RM_- *Itgae^low^* clusters. **c,** Differential distribution of cells across tissues. **d,** Kernel-density map of gRNA⁺ cells in UMAP space. **e,** Cluster distribution for each TF gRNA; TEX_term_ single-state TFs in bold. Data represent three replicates; mean ± s.e.m. Statistical analysis: two-way ANOVA with Fisher’s LSD versus gScramble. *\*\*\*\*P* < 0.0001, *\*\*\*P* < 0.001, *\*\*P* < 0.01, *\*P* < 0.05; details in Supplementary Table 5. **f,** Normalized expression of T_RM_ “up” and “down” gene-signatures^9,35^ in each KO versus control. **g-h,** Phenotypic validation: **(g)**, ratio of gRNA^+^ cells in SI to spleen and **(h)**, frequency of splenic T_CM_ (CD62L^+^ KLRG1^-^) and T_EM_ (CD62L^-^ KLRG1^+^) cells. **i- j,** T_RM_ single-state TF, *Klf6* OE enhances T_RM_ formation. P14^+^ CD8^+^ T cells transduced with *Klf6* or empty-vector were co-transferred (∼1:1) into LCMV-Armstrong-infected mice. **i,** Representative plots pre- and post-transfer. **j,** Quantification of donor CD69^+^ CD103^+^ T_RM_ cells in the SI. **k**. Quantification of the frequency of TEX_term_ cells. Statistical tests: ordinary one-way ANOVA with Dunnett’s multiple comparison versus gScramble (**g-h**), paired t-tests (**i-j**). n ≥ 4 (**g-h**) or n ≥ 6 (**i-k**) from ≥ 2 biological replicates. Data mean ± s.e.m. *\*\*\*P*<0.001, *\*\* P* <0.01, *\* P* <0.05

Cas9^+^ P14 CD8^+^ T cells were transduced with this library and transferred into mice infected with **LCMV-Clone 13**, a model of **chronic infection** and CD8^+^ T cell exhaustion (*note, recipient mice also expressed Cas9 to prevent rejection of donor cells*). Droplet-based sequencing was performed 18+ days post-transfer to assess sgRNA and transcriptomes of each spleen-derived donor Cas9^+^ P14 CD8^+^ T (Fig. 3a), analyzing 17,257 cells with unique gRNA expression.

To determine which TFs impaired TEX_term_ cell differentiation, we first employed uniform manifold approximation and projection (UMAP). Four primary clusters were identified: TEX_prog_, TEX_eff_, and TEX_term_ cells and those in cell cycle (Fig. 3b; Extended Data Fig. 6b,c). All clusters expressed *Tox* and *Pdcd1*, key exhaustion markers, and TEX_prog_ cells were identified by *Tcf7*, *Slamf6*, and *Sell* expression. TEX_eff_ cells expressed effector markers, including *Cx3cr1*, *Klrd1*, *Klrk1*, *Klf2*, and *Zeb2*^2^, while TEX_term_ cells expressed high inhibitory receptors and well-established exhaustion markers like *Cd101*, *Cd7*, *Cd38*, *Cd39*, *Cxcr6*, and *Nr4a2*. The cell cycle cluster was noted for its expression of *Birc5, Mki67*, *Stmn1*, and *Tuba1b*.

Next, we evaluated the impact of individual TF depletion by analyzing the distribution of gRNA^+^ cells across exhaustion states (Fig. 3c,d). CRISPR KO of the majority of the 19 TEX_term_ driving TFs led to a reduction in TEX_term_ cell frequency. Notably, KOs of multi-state TFs such as *Hic1*, *Stat3*, *Prdm1*, and *Ikzf3* (which encodes AIOLOS) significantly reduced TEX_term_ differentiation by 92% to 89%. Depletion of novel TEX_term_ single-state TFs—including *Zfp324*, *Zscan20*, and *Jdp2*—significantly prevented TEX_term_ differentiation by 78%, 54%, and 43%, respectively (Fig. 3d, bold). Other novel candidates such as *Etv5*, *Arid3a*, *Zfp410*, *Foxd2*, and *Prdm4* also reduced TEX_term_ representation by 25–40%, though some did not reach statistical significance. This Perturb-seq analysis highlights the platform’s ability to identify TFs that regulate the TEX_term_ state, with most tested TFs influencing exhaustion to varying degrees.

To further assess how TEX_term_-driving TF KOs affect CD8^+^ T cell exhaustion, we used flow cytometry and scRNA-seq to analyze TF KO cells during LCMV-Clone 13 infection (Fig. 3e-i; Extended Data Fig. 6). We tested six TF KOs, including known control (*Prdm1*) and five newly identified TFs (*Zscan20*, *Jdp2*, *Zfp324*, *Stat3*, *Hic1*) that impaired TEX_term_ state differentiation in Perturb-seq. Disrupting these TFs reduced TEX_term_ cell (PD1^+^ CX3CR1^-^ SLAMF6^-^) frequency by ∼50% (Fig. 3e, f) and decreased expression of inhibitory receptors such as CD101, CD39, and CD38 (Extended Data Fig. 6d,e). All 19 TEX_term_-TF KOs exhibited a marked decrease in TEX_term_-signature genes^47^, including *Cd7*, *Cxcr6*, *Nr4a2*, and *Entpd1*(Fig. 3g).

Lastly, the TEX_term_-driving TF KOs were grouped based on their effects on TEX_prog_ (PD1^+^ CX3CR1^-^ SLAMF6^+^, Fig. 3h) or TEX_eff_ (PD1^+^ CX3CR1^+^, Fig. 3i) state differentiation. Loss of *Prdm1* and *Stat3* markedly increased the frequency of TEX_prog_ cells and upregulated TEX_prog_ signature genes (Fig. 3g, h) whereas loss of *Hic1*, *Zscan20*, *Zfp324*, or *Jdp2* primarily expanded the TEX_eff_ cell population and effector signature genes (Fig. 3g, i, j). Deletion of the *Zscan20* and *Jdp2* significantly enhanced effector cytokine production (e.g., IFNγ and TNFα) and reduced viral loads in recipient mice (Fig. 3k, l).

## Deleting TEX_term_ TFs Preserves T_RM_ Fate

A major goal of this work was to identify TFs that selectively repress TEX_term_ cell differentiation without affecting T_RM_ differentiation, thereby enabling more precise programming of CD8^+^ T cell states. Since nearly all the predicted TEX_term_ single-state TFs impaired TEX_term_ differentiation to some degree (Fig. 3), the next step was to evaluate their effects on T_RM_ differentiation to confirm their selective activity. We used the same Perturb-seq library as before, but this time included only the eight TEX_term_ single- state TFs and seven multi-state TFs that impaired TEX_term_ state development by more than 25% in chronic LCMV infection (Fig. 3d). To assess their impact on memory CD8^+^ T cell development, we isolated RV-transduced Cas9^+^ P14 CD8^+^ T cells from the spleen and small intestine (SI) of mice 18 days after **acute LCMV-Armstrong** infection. We then analyzed 15,211 cells using scRNA-seq to determine how these perturbations impacted the formation of intestinal T_RM_ cells, as well as circulating splenic T_CM_ and T_EM_ cells (Fig. 4a).

The UMAP analysis identified four primary clusters containing cells with features of T_CM_ (*Il7r*, *Tcf7, Sell,* and *S1pr1),* T_EM_ (*Cx3cr1*, *Klrg1,* and *Klf2)*, and T_RM_ cells (*Cd69, Cd160, and Itgae* (encoding CD103)) as well as a small T_RM_ cell population with lower *Itgae*, but higher *Ifng* and *Irf1* expression designated T_RM_ *Itgae*^low^ (Fig. 4b,c; Extended Data Fig. 7a,b)^23,48^. Examination of the gRNA^+^ cells revealed that none of the eight TEX_term_ single-state TF KOs (*Zfp324*, *Irf8*, *Zfp410*, *Nfatc1*, *Zscan20*, *Jdp2*, *Arid3a*, and *Etv5)* negatively affected T_RM_ formation significantly (Fig. 4d,e, **bold** gene symbols). In fact, KO of *Etv5* tended to increase the frequency of T_RM_ cells. To evaluate the specificity of the TEX_term_ single-state TFs, we also examined the expression of the T_RM_ gene signatures^9,35^ in the entire population of gRNA^+^ cells for each TF tested (Fig. 4f). With the exception of *Etv5* and *Arid3a*, which increased T_RM_-signature gene expression (Fig. 4f, Extended Data Fig. 7c), perturbation of the TEX_term_ single-state TFs did not substantially alter T_RM_-signature gene expression. The platform also predicted novel multi-state TFs, including *Hic1* and *Gfi1*. Interestingly, disruption of these multi-state TFs significantly reduced T_RM_ cell frequency (Fig. 4e) and T_RM_-signature gene expression (Fig. 4f, Extended Data Fig. 7c), mirroring the effects of *Prdm1*, a known multi-state TF for T_RM_ and TEX_term_ cells.

To further validate the Perturb-seq data, we individually depleted the TEX_term_ single-state TFs *Zscan20* and *Jdp2* and multi-state TF *Prdm1* in Cas9^+^ P14 CD8^+^ T cells, adoptively transferred them into LCMV-Armstrong infected animals, and assessed their differentiation into T_CM_, T_EM_, and T_RM_ cells using flow cytometry (Fig. 4g,h). Deletion of *Zscan20* and *Jdp2* did not alter the formation of any memory cell subtypes, while perturbation of *Prdm1* significantly reduced T_RM_ and increased T_CM_ formation, as expected. Altogether, this multi-omics pipeline predicted TEX_term_ single-state TFs that drive TEX_term_ differentiation without affecting with T_RM_ cell formation and multi-state TFs that influence both cell states. These results demonstrate the accuracy and predictive power of our approach for pinpointing single-state and multi-state TFs.

## *Klf6* OE drives T_RM_ expansion without inducing exhaustion

To further demonstrate the utility of our cell-state selective TF identification pipeline in discovering novel T_RM_-associated TFs, we evaluated *Klf6*, which was identified through our Taiji analysis as a T_RM_ single-state TF (Fig. 1g). We hypothesized whether overexpressing *Klf6* (*Klf6*-OE) would enhance T_RM_ formation during acute viral infection without worsen terminal exhaustion in chronic infection. Our results confirmed this hypothesis. When empty-vector control and *Klf6-*OE P14 CD8^+^ T cells were co-transferred, *Klf6-*OE cells robustly outcompeted control cells, resulting in a 15-fold enrichment in the SI compared to controls (Fig. 4i). Furthermore, there were ∼42 times more CD69^+^ CD103^+^double-positive T_RM_-like cells in *Klf6-*OE than in control donor cells, indicating *Klf6*-OE dramatically increased T_RM_ development in the SI (Fig. 4j). Importantly, *Klf6-*OE did not increase terminal exhaustion during chronic infection (Fig. 4k, Extended Data Fig. 7d). This work not only identified KLF6 as a new T_RM_- driving TF but also confirmed its selectivity.

## Novel TEX_term_-TF loss improves tumor control

This platform predicted cell-state-selective TF activity and identified TEX_term_ single-state TFs as targets for engineering T cells that resist exhaustion yet retain effector and memory functions—offering new strategies to improve immunotherapy efficacy. Given that T_RM_ cells are associated with better clinical outcomes in solid tumors^9–12^, we hypothesized that KO of exhaustion-selective TFs like *Zscan20* could be more effective than targeting T_RM_ and TEX_term_ multi-state TFs like *Hic1*. Utilizing an ACT model, TF gRNA RV-transduced Cas9^+^ P14 CD8^+^ T cells were transferred into mice with established melanoma tumors expressing GP33-41 (Fig. 5a). Unlike depletion of the multi-state TF *Hic1*, depleting the TEX_term_ single-state TF, *Zscan20,* resulted in improved tumor control (Fig. 5b,c). Moreover, *Zscan20* gRNA^+^ cells more readily formed TEX_prog_ cells rather than TIM3^+^ CD39^+^ TEX_term_ cells (Fig. 5d,e). To control for inter-mouse variability in antigen load, we co-transferred *Zscan20* or *Hic1* KO with control P14 CD8^+^ T cells into the same B16-GP33 tumor-bearing mice (Fig. 5f–j; Extended Data Fig. 8). Both KOs significantly increased the frequency of PD1^+^ SLAMF6^+^ TIM3^-^ and decreased the frequency of TIM3^+^ exhausted cells and TEX_term_ cell state (PD1^+^ SLAMF6^-^ CX3CR1^-^) compared to controls (Extended Data Fig. 8), consistent with their predicted activity in TEX_term_ cells (Fig. 1f, h). However, *Zscan20* KO robustly enhanced effector marker expression (CX3CR1), granzyme B, and cytokine production in TILs, whereas *Hic1* KO did not appear to improve effector function to the same degree (Fig. 5g-j). Thus, despite their similar effects on suppressing TEX_term_ cell differentiation in tumors, differences in their ability to promote functional effector-like states may underlie the differential tumor control observed (Fig. 5b,c). Given that *Hic1* functions as a multi-state TF while *Zscan20* as a single-state TF, these findings support the general rationale for targeting state-specific TFs to enable more selective programming of T cell differentiation.

**Fig. 5.**
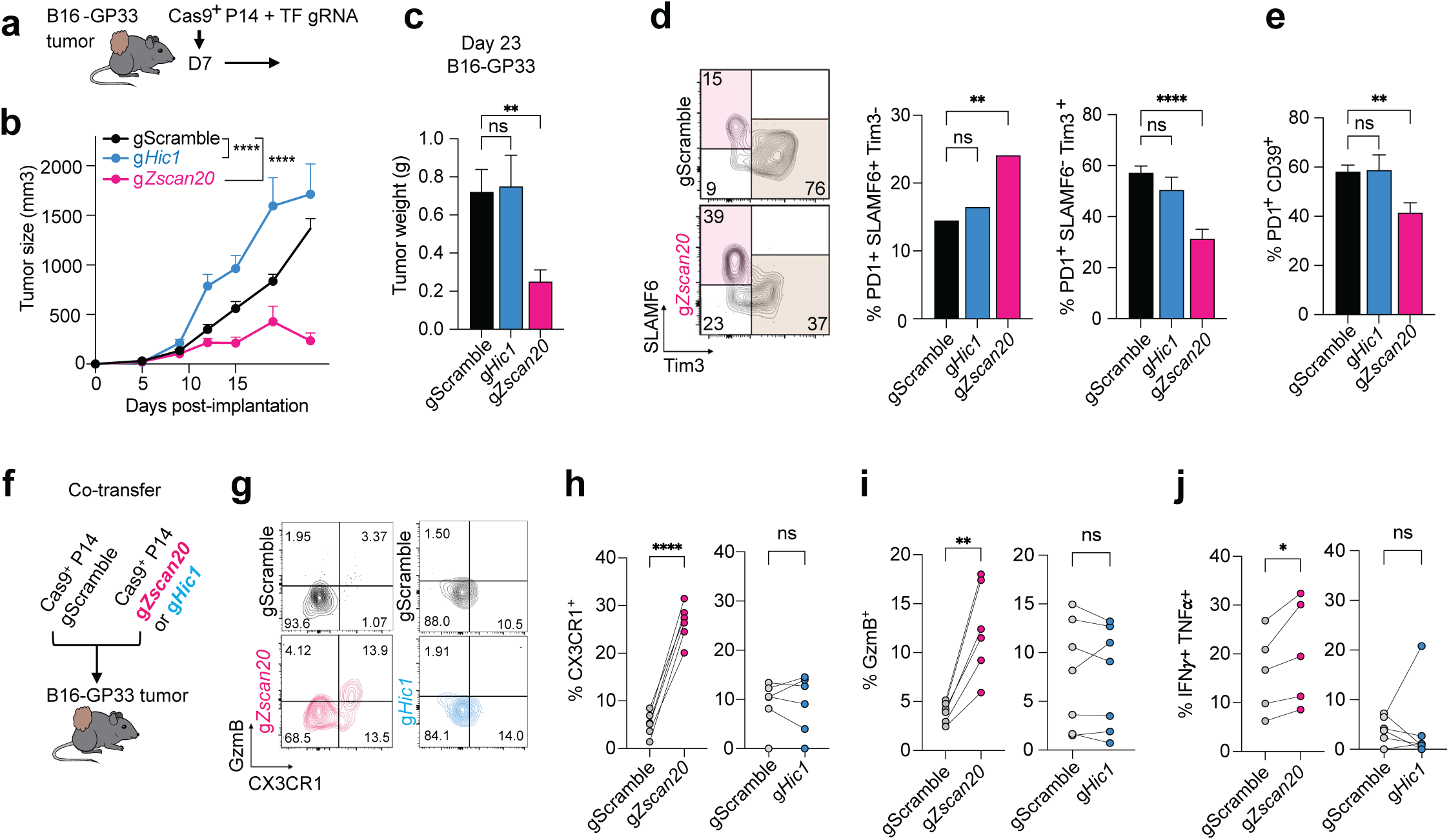
| Distinct roles of TEX_term_ single-state TF *Zscan20* and multi-state TF *Hic1* in tumor control. **a**, Experimental design: adoptive transfer of P14 CD8⁺ T cells with CRISPR KOs of either *Zscan20* (TEX_term_ single-state TF) or a *Hic1* (multi-state TF active in TEX_term_ and T_RM_) into B16-GP33 melanoma-bearing mice. **b-c**, Tumor volumes **(b)** and terminal weights **(c)** at day 23. **d**, Representative flow plots of PD1⁺ P14 cells stained for SLAMF6 and TIM3, with quantification of TEX_prog_ (PD1⁺ SLAMF6⁺ TIM3⁻) and exhausted (PD1⁺ SLAMF6⁻ TIM3⁺) subsets. **e**, Frequency of PD1^+^ CD39 ^+^ double-positive cells. **f-j**, Co-transfer design **(f)** mixing *Zscan20*- or *Hic1*-KO Cas9^+^ P14 cells with scramble-control cells before transfer. **g**, Representative plots of effector marker CX3CR1^+^ and cytotoxic cytokine GzmB in TF KO versus control. **h-j**, Quantification of CX3CR1^+^ (**h**), GzmB^+^ (**i**), and polyfunctional IFNγ^+^ TNFα^+^ (**j**) cells in *Zscan20* (left) or *Hic1* (right) KOs. Data represent n ≥ 8 (**b–e**) and n ≥ 6 (**g–j**) from ≥ 2 biological replicates. Statistical analysis: two-way ANOVA with Tukey’s multiple comparisons (**b**), one-way ANOVA with Dunnett’s test versus gScramble (**c**), paired t-tests (**g- j**). Data mean ± s.e.m. \*\*\*\**P* < 0.0001, \*\*\**P* < 0.001, \*\**P* < 0.01, \**P* < 0.05.

## State-selective TFs conserved across species

To evaluate the relevance of our murine findings in human T cells—particularly for applications in immunotherapy—we conducted cross-species validation using publicly available single-cell multi-omics and scRNA-seq datasets from human tumor-infiltrating CD8^+^ T cells (Fig. 6a; Extended Data Fig. 9a; Supplementary Table 1). Leveraging the Taiji TF analysis platform, we mapped murine TEX_term_- and T_RM_-associated TFs onto a curated human pan-cancer CD8^+^ T cell atlas encompassing six tumor types^49–56^(GBM, HNSCC, BCC, HCC, RCC, and ccRCC). Human CD8^+^ T cells were clustered into heterogeneous cell states, including T_RM_ and TEX_term_ clusters (Fig. 6a; Extended Data Fig. 9b). Taiji analysis revealed strong cross-species conservation: TEX_term_ TFs such as *JDP2*, *ZNF410*, and *FOXD2* exhibited higher activity in TEX_term_ clusters than in T_RM_-like cells (Fig. 6b). Of 34 murine TEX_term_- single-state TFs, 19 showed conserved activity patterns in human TEX_term_ cells. Similarly, T_RM_-specific TF (e.g. *NR4A1*, *KLF6*, and *FOSB*) displayed enriched activity in the human T_RM_ cluster (Extended Data Fig. 9c). Furthermore, 22 of the 30 murine TFs active in both TEX_term_ and T_RM_ states showed similar activity profiles in human datasets (Extended Data Fig. 9d). A few TFs—such as ZSCAN20—could not be assessed in the Taiji analysis due to missing DNA-binding motifs, but comparative RNA profiling across 15 tumor types supported their relevance, with 24 of 34 murine 24 of 34 murine TEX_term_ single- state TFs, including *ZSCAN20* and *JDP2*, showing higher expression in human TEX cells.

**Fig. 6.**
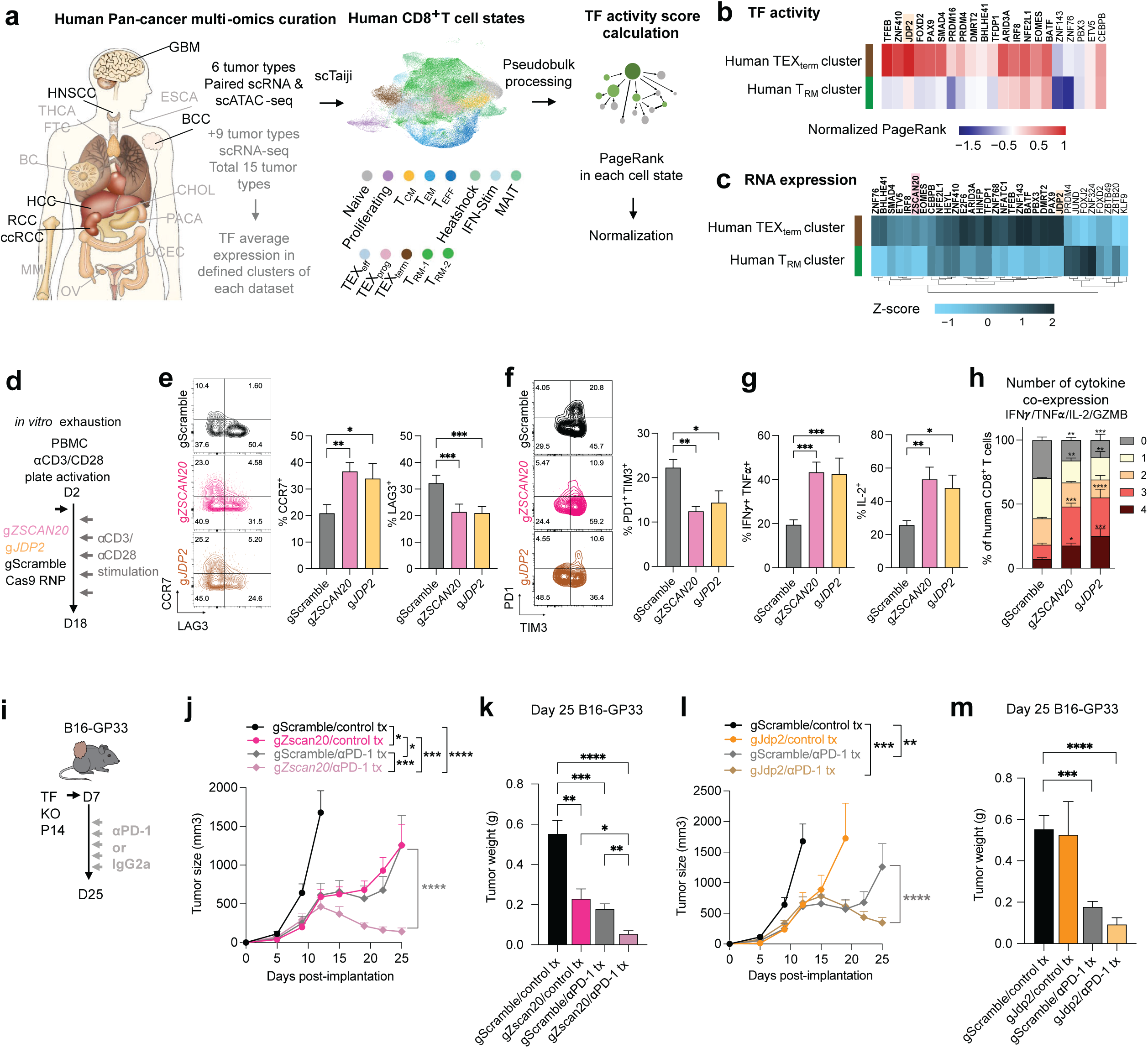
| Targeting conserved TEX_term_ single-state TFs enhances human T cell effector function and ICB response **a,** Human pan-cancer single-cell multi-omics and scRNA-seq datasets^49–56^ were integrated to evaluate TF expression and activity across CD8⁺ T cell states using the scTaiji algorithm. **b**, Paired scRNA-seq and scATAC-seq data were used to construct regulatory networks and compute PageRank TF activity scores. Shown are normalized scores of TEX_term_ single-state TFs (Fig. 1f) with conserved DNA-binding motifs in the human genome. **c,** mRNA expression of TEX_term_ TFs across TEX_term_ and TRM clusters in pan-cancer datasets; TFs with cross-species conservation in bold. **d,** Experimental design for human PBMC TF KOs. *ZSCAN20*- or *JDP2*-KO CD8⁺ T cells were stimulated with anti-CD3/CD28 beads for 18 days to model chronic activation. **e-f,** Flow cytometry analysis of CCR7 (memory/stem-like) and inhibitory receptors LAG3, PD1, TIM3 in KO versus control cells. **g,** Frequency of IFNγ^+^ TNFα^+^ and and IL-2 cells. **h,** Polyfunctionality analysis of cytokine-producing cells. **i,** Schematic of adoptive transfer and anti–PD-1 therapy to test synergy with TEXterm TF KO. Cas9⁺ P14 cells (with or without TF KO) were transferred into B16-GP33 tumor-bearing mice and treated with anti–PD-1 or isotype IgG2a. **j–m,** Tumor growth and weights of *Zscan20* (**j-k**) and *Jdp2* (**l–m)** KO versus controls. Data: mean ± s.e.m.; n ≥ 6 from ≥2 biological replicates. One-way ANOVA with Dunnett’s test (**e-g**); two- way ANOVA with Dunnett’s (**h**) or Tukey’s (**j,l**), and one-way ANOVA with Tukey’s (**k, m**). \*\*\*\**P* < 0.0001, \*\*\**P* < 0.001, \*\**P* < 0.01, \**P* < 0.05.

Given these correlations between the species, we perturbed *ZSCAN20* and *JDP2* to assess the relevance of TEX_term_ single-state TFs in human T cells (Extended Data Fig. 10a,b). Following repeated CD3/CD28 stimulation over 18 days to simulate chronic activation (Fig. 6d), *ZSCAN20*- or *JDP2*-deficient CD8^+^ T cells exhibited increased expression of CCR7 (naïve/stem cell memory/T_CM_ marker) and decreased levels of inhibitory receptors, including LAG3, PD1, and TIM3 (Fig. 6e, f). These KO cells also produced higher levels of effector cytokines and Granzyme B (Fig. 6g, h), indicating that ZSCAN20 and JDP2 contribute to exhaustion-associated features in human CD8^+^ T cells.

### ICB Synergy with *Zscan20* and *Jdp2* KOs

Tumors with high TEX_term_ cell infiltration often exhibit poor responses to immune checkpoint blockade therapy^17^. We hypothesized that targeting TEX_term_ single-state TFs could enhance ICB efficacy. Among the TEX_term_-associated TFs, *Zscan20* and *Jdp2* were prioritized for their conservation and functional relevance in human T cells (Fig. 6b-h). To test synergy with ICB, treatment began one day after adoptive transfer of TF-depleted P14 CD8^+^ T cells (Fig. 6i). The combination of *Zscan20* or *Jdp2* KO with anti-PD-1 therapy significantly reduced tumor burden (Fig. 6j–m) and improved survival (Extended Data Fig. 10c,d). These findings suggest that selectively disrupting TEX_term_ single-state TF represents a promising strategy to enhance T cell therapy by minimizing dysfunctional states while preserving beneficial T cell phenotypes. Overall, our cross-species multi-omics and functional perturbation approach underscores the translational potential of Taiji-identified TFs for improving ACT.

## Discussion

Our study introduces a powerful platform for identifying TFs pivotal in guiding specific CD8^+^ T cell state differentiation during viral infections and tumor progression. Leveraging our comprehensive transcriptional/epigenetic atlas from nine distinct CD8^+^ T cell states, we developed a detailed map of TF activity, creating a unique TF fingerprint for each context. Additionally, we developed TaijiChat, a web interface for natural language queries of our datasets and literature (Supplementary Method).

Focusing on two critical cell states TEX_term_ (dysfunctional terminally exhausted) and T_RM_ (functional, tissue-resident memory) T cells, we examined similarities and differences of TF activity and their networks in both states and engineered T cells to resist exhaustion while retaining functionality of T_RM_ cells. Using *in vivo* Perturb-seq, we validated TF activity for TEX_term_ and T_RM_ cells in both acute and chronic infection models. While recent CRISPR screenings in CD8^+^ T cells have identified important TFs in cytotoxicity, memory formation^41–43^, cell enrichment^57^, and exhaustion^58^, a systematic and context-dependent understanding of TF roles across multiple contexts has been lacking. Our study addresses this gap by generating an accurate catalogue of CD8^+^ T cell state-defining TFs, enabling cost-effective validation of predicted TF activity and selectivity via Perturb-seq. Furthermore, our study offers broader and new insight into context-dependent TF regulation. Previously, differential TF cooperation in different contexts was reported^26,43,44^. We extend this by analyzing global TF associations across cell states, revealing how TF communities regulate T cell-specific pathways, including protein catabolism in T cell exhaustion, which aligns with prior research on protein homeostasis^46,59,60^. These TF networks reveal how various cellular processes are differentially controlled between T_RM_ and TEX_term_ cells, providing rationale for their different functional capabilities within tissues.

One of the key outcomes of this study was the identification of novel TFs, including ZSCAN20 and JDP2 as TEX_term_ single-state TFs, KLF6 as a T_RM_ single-state TF, and newly uncovered roles for multi-state TFs such as HIC1 and GFI1. Perturbing TEX_term_ single-state TFs not only prevented T cell exhaustion but also preserved the ability of these cells to differentiate into effector and memory states. This led to significant improvements in tumor control.

To evaluate the clinical importance of the newly discovered TFs and the catalog of TFs with TEX_term_ and T_RM_ selectivity, we confirmed cross-species conservation of a significant number of TFs using Taiji analysis of a human pan-cancer multi-omics atlas, along with comparative expression analysis across pan-cancer single-cell RNA-seq datasets. Furthermore, we demonstrated enhanced human T cell function following perturbation of the TEX_term_ single-state TFs *ZSCAN20* and *JDP2*. Depletion of these TFs shows synergistic effects with ICB therapy, leading to significant tumor regression. These findings highlight a promising strategy for enhancing antitumor immunity through precise cell-state programming.

Our TF atlas-guided platform can offer optimized ‘TF recipes’ for cell programming with increased precision, robustness, and durability. Future strategies may combine overexpression of T_RM_ -promoting TFs like KLF6 with depletion of TEX_term_ TFs. Such recipes can be refined with AI models. In summary, while our study focuses on CD8^+^ TEX_term_ and T_RM_ cell differentiation, the pipeline for identifying single- state TFs and ‘TF recipes’ can be adapted for other cell types, expanding cell therapy applications.

## Acknowledgments

We thank members of the Chung, Kaech, and Wang laboratory for their advice and assistance with experiments. We appreciate UNC Lineberger Preclinical Research Unit at the University of North Carolina at Chapel Hill assisted animal experiments which is supported in part by an NCI Center Core Support Grant (CA16086) to the UNC Lineberger Comprehensive Cancer Center. This work was supported by NIH grants to S.M.K. R37AI066232, R01AI123864, R21AI151986, and R01CA240909, to W.W. R01AI150282, R01HG009626 (in part) to H.K.C. K01EB034321, to J.J.M. R01AI177864 to

J.E.T. R01CA248359 and R01CA244361, to D.C.H. AI151123, to Y.W. EB029122 and GM140929, and to L.L. K01EB035649. H.K.C. is a Damon Runyon Fellow supported by Damon Runyon Cancer Research Foundation grant [DRG-2374-19].

## Author contributions

H.K.C. designed and performed experiments. A.B., B.M.R., F.X., Y.L., K.Y., D.J., M.S., J.G., S.M., E.C., B.M. D.C., B.C., Q.Y., F.H., T.H.M., S.K.V., V.T., U.H.C., G.D., B.S., Y.W., L.L., P.H., and J.H. assist with experiments. H.K.C., C.L., A.J., N.J.B., M.S. analyzed the sequencing and bioinformatics data. J.W., B.M., M.S., Y.H., L.H., U.H.C., J.M., contribute to bioinformatics data analysis. Z.W., A.W. contributed to TaijiChat and portal generation. J.H. and D.H. provided resources and training for ATAC- sequencing. B.P.R. and J.E.T. performed proteasome activity experiments. H.K.C., S.M.K., W.W., Y.W., L.L., B.S., G.P., J.E.T., J.J.M., D.C.H, provided scientific input and acquired funding. H.K.C, and C.L. wrote the original manuscript. H.K.C., W.W., and S.M.K. reviewed and edited the paper. H.K.C., S.M.K., and W.W. conceptualized, designed, and supervised the study.

## Additional information

Supplementary Information is available for this paper. Correspondence and requests for materials should be addressed to H. Kay Chung, Susan M. Kaech, and Wei Wang.

## Methods

### Dataset acquisition for murine CD8^+^ T cell state multiomics atlas

CD8^+^ T cell samples were collected from ten datasets, including those generated in this study (Extended Data Fig. 1e). In total, we analyzed 121 experiments, comprising 52 ATAC-seq and 69 RNA-seq datasets, which were integrated to generate paired samples and served as input for the Taiji pipeline. The samples encompassed nine distinct CD8^+^ T cell subtypes: naive cells (Naive), terminal effector cells (TE), memory precursor cells (MP), tissue-resident memory cells (T_RM_), effector memory cells (T_EM_), central memory cells (T_CM_), progenitor exhausted cells (TEX_prog_), intermediate exhausted cells (TEX_eff_) and terminal exhausted cells (TEX_term_). Cell states were defined based on established surface marker combinations and LCMV (lymphocytic choriomeningitis virus)-specific tetramers, including IL7R, KLRG1, PD1, SLAMF6, CD101, Tim3, CD69, CD103, H2-Db LCMV gp 33-41 and H2-Db LCMV gp 276-286 or congenic markers for P14 (T cell receptor (TCR) specific for the LCMV GP33–41 peptide CD8^+^ T cells), in the context of either acute (LCMV-Armstrong) or chronic (LCMV-Clone 13) infection models. A complete summary of dataset sources, accession numbers, infection conditions, and corresponding cell state definitions (sorting gates) is provided in Supplementary Table 1 and Extended Data Fig. 1e.

### TF regulatory networks construction and visualization

To perform integrative analysis of RNA-seq and ATAC-seq data, we developed Taiji v2.0. which allows visualization of several downstream analysis-TF wave, TF-TF association, TF community analysis. Epitensor was used for the prediction of chromatin interactions. Putative TF binding motifs were curated from the latest CIS-BP database^62^. In this analysis, 695 TFs were identified as having binding sites centered around ATAC-seq peak summits. The average number of nodes (genes) and edges (interactions) of the genetic regulatory networks across CD8^+^ T cell states were 15,845 and 1,325,694, respectively, including 695 (4.38%) TF nodes. On average, each TF regulated 1,907 genes, and each gene was regulated by 22 TFs.

### Identification of single-state/multi-state TFs

We first identified universal TFs with mean PageRank across 9 cell states ranked as top 10% and Coefficients of variation (CV) less than 0.5. In total, 54 universal TFs were identified and shown in Supplementary Table 1. The remaining 641 TFs were candidate TFs for single-state TFs. To identify single-state TFs, we divided the samples into two groups: target group and background group. Target group included all the samples belonging to the cell state of interest and the background group comprised the remaining samples. We then performed the normality test using Shapiro-Wilk’s method to determine whether the two groups were normally distributed, and we found that the PageRank scores of most samples (90%) follow a log-normal distribution. Based on the log-normality assumption, an unpaired t-test was used to calculate the P-value. A P-value cutoff of 0.05 and log2 fold change cutoff of 0.5 were used for calling lineage-specific TFs In total, 255 specific TFs were identified (Supplementary Table 2). Depending on whether the TF appeared in multiple cell states, they could be further divided into multi-state TFs (Supplementary Table 2, Fig 1e) and 136 single-state exclusive TFs (Supplementary Table 2, Fig. 1d). Out of 255 single-state TFs, 84 TFs appear in TEX_term_ or T_RM_. To identify the truly distinctive TFs between TEX_term_ and T_RM_, we performed a second round of unpaired t-test only between TEX_term_ and T_RM_ (Supplementary Table 3). The same cut-offs, i.e. P-value of 0.05 and log2 fold change of 0.5, were applied to select TEX_term_ single-taskers and T_RM_ single- taskers. Thirty out of 84 TFs that did not pass the cut-off were identified as TEX_term_ and T_RM_ multi- taskers. The full workflow was summarized in Extended Data Fig. 2a.

### Identification of transcriptional waves

Combined with prior knowledge on the T cell differentiation path, TF waves are combinations of TFs which are particularly active in certain differentiation stages, revealing possible mechanisms of how TF activities are coordinated during differentiation. To be more specific, we clustered the TFs based on the normalized PageRank scores across samples. First, we performed principal component analysis (PCA) for dimensionality reduction of the TF score matrix. We retained the first 10 principal components for further clustering analysis, which explained more than 70% variance (Extended Data Fig. 3b; left panel). We used the k-means algorithm for clustering analysis. To find the optimal number of clusters and similarity metric, we performed the Silhouette analysis to evaluate the clustering quality using five distance metrics: Euclidean distance, Manhattan distance, Kendall correlation, Pearson correlation, and Spearman correlation (Extended Data Fig. 3b; right panel). Pearson correlation was the most appropriate distance metric since the average Silhouette width was the highest among all five distance metrics. Based on these analyses, we identified 7 distinct dynamic patterns of TF activity during immune cell development. We further performed functional enrichment analysis to identify GO terms for these clusters.

### TF-TF collaboration network analysis and visualization

To build the TF-TF association networks, we first defined a set of relevant TFs for each context (TEX_term_ and T_RM_) by combining cell state- important and single-state TFs, resulting in 159 TFs for TEX_term_ and 170 for T_RM_. The analysis was based on a TF-regulatee network derived from Taiji, where we first consolidated sample networks by averaging the edge weights for each TF-regulatee pair. To reduce noise, regulatees with low variation across all TFs (standard deviation ≤ 1) were removed. Subsequently, a TF-TF correlation matrix was generated by calculating the Spearman’s correlation of edge weights for each TF pair across their common regulatees. From this matrix, we constructed a graphical model using the R package “huge”^63^, which employs the Graphical Lasso algorithm and a shrunken ECDF (empirical cumulative distribution function) estimator. An edge between two TFs was established if their correlation was deemed significant by the model, controlled by a lasso penalty parameter (lambda) of 0.052. This value was chosen as it represents a local minimum on the sparsity-lambda curve, resulting in approximately 15% of TF-TF pairs being connected. To validate this method, we estimated the false discovery rate by generating a null model through random shuffling of the TF-regulatee edge weights. Applying our algorithm to this null data identified zero interactions, confirming that our approach has a very low false discovery rate.

### TF community construction and visualization

Following the construction of the TF-TF association networks, we identified functionally related TF communities within each network. We applied the Leiden algorithm^64^, using modularity as the objective function and setting the resolution parameter to 0.9, as this value achieved the highest clustering modularity in our analysis. This procedure identified five distinct communities for each context (TEX_term_ and T_RM_). The final networks, with their detected communities, were visualized using the Fruchterman-Reingold layout algorithm^65^ to spatially represent the TF-TF association structure.

### Pathway enrichment analysis

The enriched functional terms in this study were analyzed by the R package clusterProfiler 4.0.5. We used GO, KEGG, MSigDb databases for annotation. For gene set enrichment analysis (GSEA), the genes were first ranked by the mean edge weight in corresponding samples and H, C5, C6, and C7 collections from MSigDb were used for annotation. A cutoff of P-value < 0.05 was used to select the significantly enriched GO terms and KEGG pathways.

## Heuristic score calculation via the integration of TF regulatory network and Perturb-seq

We reasoned that (1) log2FC change in expression due to TF KO, and (2) TF-gene regulatory edge weights could be combined to provide heuristic scores for a TF’s regulatory effect on a target gene. For each guide RNA KO, Seurat’s FindMarkers() function was used to quantify log2FC of gene’s expression with respect to the gScramble condition. Heuristic scores were calculated for each TF-gene pair by multiplying the gene log2FC with the corresponding edge weight from the Taiji analysis. Regulatees of a TF were annotated as high-confidence if the magnitude log2FC of the regulate exceeded 0.58 (corresponding to a fold-change of 1.5 or its reciprocal) and if the edge weight belonged to the upper quantile of all edge weights attributed to the TF. The sign of the log2FC was used to determine if the TF activated or repressed each target gene.

## Human TF activity and cell-state selectivity analysis

*Taiji-based analysis of human multiomic datasets.* Transcription factor (TF) activity was inferred using the Taiji pipeline applied to matched single-cell RNA-seq (scRNA-seq) and ATAC-seq (scATAC-seq) datasets from various human cancers, including ccRCC (n=2; PRJNA768891, GSE240822), GBM (GSE240822), BCC (GSE123814, EGAS00001006141), HNSCC (GSE139324, EGAS00001006141), HCC (GSE125449, EGAS00001006141), and RCC (PMID: 30093597; EGAS00001006141). Cell types were annotated using canonical marker gene expression and categorized into six major CD8+ T cell states: T_Ex (exhausted), T_ExProg (progenitor exhausted), T_Eff (effector), T_RM (tissue-resident memory), T_CM/Naive (central memory/naive), and proliferating. PageRank scores derived from Taiji were log-transformed, averaged within each T cell category, and then standardized using z-score normalization. Results were visualized with a focus on TF activity in T_RM and T_Ex populations.

*TF expression comparison across human CD8+ T cell states.* To assess TF expression across diverse T cell states, raw count matrices from a published pan-cancer CD8+ T cell atlas (GSE156728) were reprocessed using Seurat’s standard workflow. The dataset encompassed T cells from 11 tumor types, including BC, BCL, CHOL, ESCA, FTC, MM, OV, PACA, RC, THCA, and UCEC. Cell type annotations provided by the original study were retained and mapped to the following broad categories: T_RM, T_Ex, T_EM (effector memory), T_m (memory), T_n (naive), and T_k (cycling). Seurat’s AverageExpression function was used to compute average logCPM expression for each TF in each T cell category, followed by z-score normalization. Data visualization emphasized comparisons between T_RM and T_Ex subsets.

### Mice and infections

C57BL/6/J, OT-1 (C57BL/6-Tg(TcraTcrb)1100Mjb/J), B6.Cg-*Rag2^tm^*^1^*^.1Cgn^*/J, CD45.1 (B6.SJL-*PtprcaPepcb*/BoyJ) mice were purchased from Jackson Laboratories. P14 mice (Pircher et al., 1987) mice have been previously described. Cas9 P14 mice were generated by crossing P14 mice with B6(C)-Gt(ROSA)26Sorem1.1(CAG-cas9*,-EGFP)Rsky/J (Jackson Laboratories). Animals were housed in specific-pathogen-free facilities at the Salk Institute and University and at the University of North Carolina at Chapel Hill. All animal experiments were approved by the Institutional Animal Care and Use Committee (IACUC). Mice were infected with 2x10^5^ PFU LCMV-Armstrong by intraperitoneal injection or 2x10^6^ PFU LCMV-Clone13 by retro-orbital injection under anesthesia.

### Viral titers

LCMV Fluorescence Focus Unit titration was performed seeding Vero cells at a density of 30k cells/100µl in a 96 well flat bottom plate in DMEM + 10% FBS + 2% HEPES + 1% Pen/Strep. On the next day, tissues were homogenized on ice, spun down at 1000g for 5 min at 4°C and supernatants or serum were diluted in 10-fold steps. Diluted samples were added to Vero cells and incubated at 37°C, 5% CO_2_ for ∼20h. Subsequently, inocula were aspirated and wells were incubated with 4% PFA for 30 min at RT before washing with PBS. VL-4 antibody (BioXCell) was conjugated using the Invitrogen AF488 conjugation kit and added to the wells in dilution buffer containing 3% BSA and 0.3% Triton (Thermo Fisher Scientific) in PBS. Cells were incubated at 4°C overnight before washing with PBS and counting foci under the microscope. FFU was calculated per this formula: FFU/ml=Number of plaques/ (dilution× volume of diluted virus added to the plate).

### Cell isolation

Spleens were mechanically dissociated with 1mL syringe plungers over a 70um nylon strainer. Spleens were incubated in ammonium chloride potassium (ACK) buffer for 5 minutes. For isolation of small intestinal IEL, Peyer’s patches were first removed by dissection. Intestines were longitudinally cut and then cut into 1cm pieces and washed in PBS. Pieces were incubated in 30mL HBSS with 10% FBS, 10mM HEPES, and 1mM dithioerythritol with vigorous shaking at 37C for 30 minutes. Supernatants were collected, washed, and further isolated using a 40/67% discontinuous percoll density centrifugation for 20 minutes at room temp with no brakes.

### Cell lines and *in vitro* cultures

B16-GP33 melanoma cell line were cultured in DMEM (Invitrogen) with 10% fetal bovine serum, 1% penicillin-streptomycin and 250 μg/ml G418 (Invitrogen #10131027). MCA-205 (Sigma) tumor line was maintained in RPMI supplemented with 10% FBS, 300 mg/L L-glutamine, 100 units/mL penicillin, 100 μg/mL streptomycin, 1mM sodium pyruvate, 100μM NEAA, 1mM HEPES, 55μM 2-mercaptoethanol, and 0.2% Plasmocin mycoplasma prophylactic (InvivoGen). All the tumor cell lines were used for experiments when in the exponential growth phase. For *in vitro* T cell culture, splenocytes were activated in RPMI 1640 medium (Invitrogen) containing 10% fetal bovine serum and 1% penicillin-streptomycin, 2mM L-glutamine, 0.1 mg/ml GP33, beta-mercaptoethanol 50 mM and 10 U/ml IL-2.

### Tumor engraftment and treatment of tumor-bearing mice (Fig. 5, 6)

3×10^5^ B16-GP33 (Fig. 5), 5×10^5^ B16-GP33 tumor cells (Fig. 6) were injected subcutaneously in 100 μl PBS. Around 0.5-1x10^6^ Cas9^+^ P14 T cells with CD45.1 markers were transferred to tumor on day 7 without pre-radiation of tumor-bearing mice. Tumors were measured every 2-3 days post-tumor engraftment for indicated treatments and calculated. Tumor volume was calculated by volume = (length × width^2^)/2. For antibody-based treatment, tumor-bearing mice were treated with anti-PD1 antibody (200 µg per injection, clone RMP1-14, BioXcell) twice per week from day 7 post-tumor implantation. Tumor growth was measured twice per week with calipers. Survival events were recorded each time a mouse reached the endpoint (tumor volume ≥1,500 mm^3^). Tumor weights were measured on day 23 for Fig. 5 and on day 25 for Fig. 6. All experiments were conducted according to the Salk Institute Animal Care and Use Committee.

### Tumor digestion and cell isolation for Fig. 5

Tumors were minced into small pieces in RPMI containing 2% FBS, DNase I (0.5 µg/ml, Sigma-Aldrich), and collagenase (0.5 mg/ml, Sigma-Aldrich) and kept for digestion for 30 min at 37°C, followed by filtration with 70 µm cell strainers (VWR). Filtered cells were incubated with ACK lysis buffer (Invitrogen) to lyse red blood cells, mixed with excessive RPMI 1640 medium (Invitrogen) containing 10% fetal bovine serum and 1% penicillin-streptomycin, and centrifuged at 400g for 5 min to obtain a single-cell suspension.

### Proteasome activity analysis

For experiments involving the Proteasome Activity Probe (R&D systems), cells of interest were incubated with the probe at concentration of 2.5mM for 2 hours at 37C in PBS. Samples were washed and then stained with Zombie NIR viability dye (Biolegend) in PBS at 4C for 15 minutes. Samples were then stained with some variation of the following antibodies for 30 minutes in FACS Buffer on ice: CD45-BV510 (BD Biosciences), CD45.2-BV510 (Biolegend), CD45.1- PE-Cy7 (Invitrogen), CD4-APC Fire 810 (Biolegend), CD11b-Alexa Fluor 532 (Invitrogen), CD8-Spark NIR 685 (Biolegend), CD44-Brilliant Violet 785 (Biolegend), CD62L-BV421 (BD Biosciences), PD1-BB700 (BD Biosciences), TIM-3-BV711 (Biolegend), LAG-3-APC-eFluor 780 (Invitrogen), SlamF6-APC (Invitrogen), CD39-Superbright 436 (Invitrogen), CX3CR1-PE/Fire 640 (Biolegend), CD69-PE-Cy5 (Biolegend), GITR-BV650 (BD Biosciences), CD27-BV750 (BD Biosciences). Samples were collected on a Cytek Northern Lights and analyzed via Cytek SpectraFlo software.

### Tumor experiment for proteasome assay (Fig. 2)

MCA-205 fibrosarcomas (2.5 x 10^5^) were established via subcutaneous injection into the right flank of C57BL/6 mice. After 12-14 days of tumor growth, spleens, draining lymph nodes, and tumors from groups of mice were harvested and tumors were processed using the Mouse Tumor Dissociation Kit and gentleMACS dissociator (Miltenyi Biotec) according to manufacturer’s protocol. For purification experiments, samples were pre-enriched using the EasySep Mouse CD8^+^ T Cell Isolation Kit (Stemcell Technologies) according to manufacturer’s protocol and stained with Live-or-Dye PE Fixable Viability Stain (Biotium) and CD8a-APC (Invitrogen) and live CD8^+^ cells were sorted using the FACSAria II cell sorter. CD8^+^ spleen and pooled TIL samples were washed in PBS and frozen for RNA-seq analysis. For adoptive cellular therapy experiments, B16- GP33 melanomas were established subcutaneously by injecting 5.0×10^5^ cells into the right flank of CD45.1 mice and tumor-bearing hosts were irradiated with 5 Gy 24 hours prior to T cell transfer. In contrast, mice used in the experiments shown in Fig. 5 and Fig. 6 were not irradiated prior to T cell transfer. After 7 days of tumor growth, 1.5×10^6^ CD45.2 OT-1 T cells and 1.5×10^6^ CD45.1/CD45.2 P14 T cells were infused in 100 μL PBS via tail vein into tumor bearing mice. Tumors were harvested 14 days after adoptive cell transfer and CD8 TILs were analyzed for proteasome activity. All experiments were conducted in accordance with the guidelines of the University of North Carolina at Chapel Hill Animal Care and Use Committee.

### Proteasome^high^/Proteasome^low^ T cell adoptive transfer experiment (Fig. 2 and Extended Data Fig. 5c)

For the adoptive transfer experiment involving proteasome^high^ and proteasome^low^ tumor-specific OT-1 T cells (Fig. 2l), whole splenocytes from OT-1 mice were activated with 1 μg/mL OVA_257–264 peptide and expanded for 7 days in the presence of 200 U/mL rhIL-2 (NCI). On day 7, OT-1 cells were FACS-sorted based on proteasome activity to isolate proteasome^high^ and proteasome^low^ OT-1 populations. A total of 2.5 × 10^5^ sorted OT-1 cells were injected into C57BL/6 mice bearing B16F1- OVA melanomas. Tumors were established by subcutaneous injection of 3 × 10^5^ B16F1-OVA cells into the right flank 7 days prior to T cell transfer. Recipient mice were preconditioned with 5 Gy total body irradiation 24 hours before adoptive transfer. Tumor growth was measured every other day with calipers. For Extended Data Fig. 5c, MCA-205 fibrosarcomas (2.5 x 10^5^) were established via subcutaneous injection into the right flank of C57BL/6 mice. After 14 days of tumor growth, Live CD45^+^ CD8^+^ CD44^+^ PD1^+^ T cells were sorted from tumors based on proteasome activity (high vs low) using the FACSAria II cell sorter. 2.5 x 10^4^ cells were then injected into the 2 day MCA-205 bearing RAG2-/- hosts (n=5 per group) and tumor growth was monitored every other day starting on day 4. All experiments were conducted in accordance with the guidelines of the University of North Carolina at Chapel Hill Animal Care and Use Committee.

### Retrovirus transduction and adoptive transfer

For overexpression of the gRNA retrovirus vector, 293T cells were transfected with the Eco-helper and MSCV gRNA vectors. 48 h and 72 h later, supernatant containing retroviral particles was ready for transduction. Donor P14 splenocytes were *in vitro* activated by 0.1 mg/ml GP33 and 10 U/ml IL-2 at 37°C for 24h, then spin-transduced (1500 g) with fresh RV supernatant from 293T cells for 90 min at 30°C in the presence of 5 μg/ml polybrene.

### CRISPR-Cas9/RNP nucleofection

Naïve CD8+ T cells were enriched from spleen using the EasySep Mouse CD8+ T cell Isolation Kit (STEMCELL Technologies). sgRNAs targeting *ZSCAN20, JDP2, Etv5, Prdm1, Hic1* genes or the mouse or human genome nontargeting scramble (control) were obtained from Synthego, IDT, and GeneScript (Supplementary Table 5). Cas9 RNP was prepared immediately before experiments by incubating 1 µl sgRNA (stock: 3 nmol in 10 µl water), 0.6 µl Cas9 nuclease (IDT, stock: 62 µM) and 3.4 µl RNase-free water at room temperature for 10min. Nucleofection of naïve CD8+ T cells was performed using Lonza (Allendale, NJ) P3 primary cell kit and program DN100 with 4D-Nucleofector (Lonza Bioscience) for mouse and EO115 for human stimulated T cells . Each nucleofection reaction consisted of approximately 5-10 × 10^6^ cells in 20 µl of nucleofection reagent and mixed with 5 µl of RNP: Cas9 complex. After electroporation, 100 µl of T cell culture media was added to the well to transfer the cells to 1.5 mL Eppendorf tube. The cells were rested at 37°C for 3 min. For *in vivo* adoptive transfer, cells were resuspended in PBS at desired concentration and adoptively transferred into recipient mice.

### CRISPR gene editing validation via Sanger sequencing

The genomic DNA (gDNA) was isolated from both the KO-induced CD8+ T cells and the control cells using the Quick-DNA MicroPrep Kit (Zymo). The gDNA concentrations were quantified using the NanoDrop One spectrophotometer (ThermoFisher Scientific). Following isolation, PCR amplification was performed with 2x Phusion Plus Green PCR Master Mix (ThermoFisher Scientific) and the respective validation primers under the following conditions: (98 °C for 5 mins; 35 x of 98 °C for 10 secs, 69 °C for 20 secs, 72 °C for 20-30 secs/kb; 72 °C for 2 mins; hold at 10 °C). The PCR products were run on a 2% agarose gel with SYBR Safe DNA Gel Stain (Invitrogen), and the appropriate bands on the gel were extracted and purified with the Gel DNA Recovery Kit (Zymo). Concentrations of the purified amplicon samples were measured and then sent off for sequencing along with the designated sequencing primers designed from Benchling’s Primer3 tool. The samples with the KOs were compared to the wildtype controls using EditCo’s Ice Analysis software, providing the indel percentages, KO score, and the indel distributions used to assess the editing efficiency. Indel% ranges from 56% to 97%, and the KO score throughout experiments ranges from 32 to 74.

### Flow cytometry, cell sorting, and antibodies

Both single cell suspensions were incubated with Fc receptor-blocking anti-CD16/32 (BioLegend) on ice for 10 min before staining. Cell suspensions were first stained with Red Dead Cell Stain Kit (ThermoFisher) for 10 min on ice. Surface proteins were then stained in FACS buffer (PBS containing 2% FBS and 0.1% sodium azide) for 30 min at 4°C. To detect cytokine production *ex-vivo*, cell suspensions were re-suspended in RPMI 1640 containing 10% FBS, stimulated by 50 ng/ml PMA and 3 μM Ionomycin in the presence 2.5 μg/ml Brefeldin A (BioLegend #420601) for 4 h at 37°C. Cells were processed for surface marker staining as described above. For intracellular cytokine staining, cells were fixed in BD Cytofix/Cytoperm (BD #554714) for 30 min at 4 °C, then washed with 1× Permeabilization buffer (Invitrogen #00-8333-56). For transcription factor staining, cells were fixed in Foxp3 / Transcription Factor Fixation/Permeabilization buffer (Invitrogen #00-5521-00) for 30 min at 4 °C, then washed with 1× Permeabilization buffer. Cells were then stained with intraceulluar antibodies for 30 min at 4 °C. Samples were processed on LSR-II flow cytometer (BD Biosciences) and data were analyzed with FlowJo V10 (TreeStar). Cells were sorted either on FACSAria™ III sorter or Fusion sorter (BD Biosciences). The following antibodies against mouse proteins were used: anti-CD8a (53-6.7), anti-PD1 (29F.1A12), anti-CX3CR1 (SA011F11), anti- SLAMF6 (13G3), anti-CD38 (90), anti-CD39 (24DMS1), anti-CD101 (Moushi101), anti-KRLG1 (2F1), anti-CD69 (H1.2F3), anti-CD103 (M290), anti-CD62L (MEL-14), anti-Tim3 (RMT3-23), anti-Ly5.1 (A20), anti-Ly5.2 (104), anti-IFN-γ (XMG1.2), anti-TNF-α (MP6-XT22). The following antibodies against human proteins were used: anti-CD8a (RPA-T8), anti-CD4 (SK3), anti-CD45RA (H100), anti-CD45RO (UCHL1), anti-CCR7 (G043H7), anti-CD62L (DREG-56), anti-CD69 (FN50), anti-CD103 (Ber-ACT8), anti-CXCR6 (K041E5), anti-PD1 (EH12.2H7), anti-CD38 (HIT2), anti-CD39 (A1), anti-LAG3 (11C3C65), anti-Tim3 (F38-2E2), anti-TIGIT (A15153G), anti-IFN-γ (4S.B3), anti-TNF-α (MAb11), anti-IL 2 (JES6-5H4), anti-GZMB (QA16A02), anti-G4S Linker (E7O2V). These antibodies were purchased from Invitrogen, Biolegend, Cell Signaling, or eBiosciences.

### *In vivo* individual TF KO phenotyping

To assess the functional impact of individual TF KOs in CD8⁺ T cells, we used Cas9-expressing P14 donor cells (LCMV-specific TCR transgenic mice, CD45.1 congenic) transduced with GFP-expressing retroviral vectors encoding individual gRNAs. Transductions were performed on the day of adoptive transfer without prior sorting. Without sorting, transduced donor cells (0.5-1 x 10^5^) were immediately transferred into congenically distinct Cas9- expressing wild-type recipient mice (CD45.2) infected one day prior with either LCMV-Clone 13 or Armstrong strains. At ≥ day 20 post-infection, spleens from Clone 13 model and spleens and small intestines from Armstrong model were harvested. Single-cell suspensions were prepared and analyzed by flow cytometry. Live, single cells were first gated on CD8⁺ cells, followed by gating on CD45.1⁺ P14 donor CD8^+^ T cells. Successfully transduced (gRNA⁺) cells were identified by GFP expression, which ranged from 10% to 70% of P14 CD8^+^ T cells across experiments. Due to variability in the number of GFP⁺ donor P14 CD8^+^ T cells obtained from different experiments, all phenotypic analyses were performed within the GFP⁺ (gRNA⁺) CD45.1⁺ CD8⁺ population. PD-1 positive and negative cells, exhaustion subsets (TEX_term_: PD-1⁺ SLAMF6⁻ CX3CR1⁻, TEX_prog_ :PD-1⁺ SLAMF6⁺ CX3CR1⁻, TEX_eff_: PD-1⁺ CX3CR1⁺) or expression of phenotypic markers was reported as a percentage within the gRNA⁺ (GFP⁺) P14 CD8^+^ T cell population to ensure consistency across samples.

### Co-transfers of control and TF KO/overexpression P14 CD8^+^ T cells in infection or tumor models

Naive CD8⁺ T cells were isolated from the spleens and lymph nodes of Cas9-expressing LCMV TCR transgenic (Cas9 P14) or P14 mice using the EasySep™ Mouse CD8⁺ T Cell Isolation Kit (STEMCELL Technologies). Purified P14 cells were activated for ∼24 hours on plates coated with goat anti-hamster IgG (ThermoFisher), followed by 1 μg/mL hamster anti-mouse CD3 and 1 μg/mL hamster anti-mouse CD28 antibodies (ThermoFisher), in complete T cell medium (RPMI-1640 supplemented with 10% FBS (HyClone), 55 μM 2-mercaptoethanol, 100 IU/mL penicillin-streptomycin, and 1% HEPES). After activation, cells were transduced with retroviruses encoding Klf6 overexpression or gRNAs targeting Hic1 or Zscan20, and cultured with 20 IU/mL IL-2, 2.5 ng/mL IL-7, and 2.5 ng/mL IL-15 (PeproTech). At 48 hours post-transduction, reporter expression was confirmed by flow cytometry. Donor cell mixes were prepared using control vs. Klf6-overexpressing cells (Fig. 4), or gScramble vs. gHic1/gZscan20 cells (Fig. 5). For LCMV infection studies, 1.5 × 10⁵ transduced P14 CD8^+^ T cells were transferred into recipient mice, followed by infection with either 2 × 10⁵ PFU LCMV-Armstrong (acute infection, intraperitoneal) or 2 × 10⁶ PFU LCMV-Clone 13 (persistent infection, intravenous). For tumor studies, 5 × 10⁵ to 1 × 10⁶ transduced T cells (gScramble vs. gTF) were transferred on day 7 after B16-GP33 tumor implantation. All experiments were conducted according to the University of North Carolina at Chapel Hill Animal Care and Use Committee.

## Perturb seq screening using the retroviral transcriptional factor library

D*ual-guide direct-capture retroviral sgRNA vector.* To generate dual-guide sgRNA vector (MSCV- hU6-mU6-SV40-EGFP), we replaced the hU6 RNA scaffold region of the previously described retroviral sgRNA vector MG-guide^66^ with an additional scaffold^67^ and murine U6 promoter.

*Dual-guide direct-capture retroviral library construction.* For the curated gene list containing 21 TFs, a total of four gRNA sequences distributed on two individual constructs were designed for each gene. To construct the library, a customized double-strand DNA fragment pool containing 80 oligonucleotides targeting those 19 TFs and 4 scramble gRNAs (each oligonucleotide contains two guides targeting the same gene) (Supplementary Table 5) was ordered from IDT. The dual-guide library was generated using an In-Fusion (Takara) reaction. In brief, the gRNA containing DNA fragment pool was combined in MG-guide vector linearized with BpiI (Thermo Fisher). The construct was then transformed into Stellar competent cells (Takara) and amplified, and the resulting intermediate, individual construct was assessed for quality using Sanger sequencing. Then, individual dual-gRNA vectors were combined. For quality control, sgRNA skewing was measured using the MAGeCKFlute^68^ to monitor how closely sgRNAs are represented in a library.

*In vivo screening.* Retrovirus was generated by co-transfecting HEK293 cells with the dual-guide, direct-capture retroviral TF library and the packaging plasmid pCL-Eco. Supernatants were collected at 48 and 72 hours post-transfection, then stored at −80 °C. Cas9-expressing P14 CD8^+^ T cells were transduced with the viral supernatant to achieve a transduction efficiency of 20–30%. To ensure sufficient representation of control cells in downstream analysis, 50% of the viral mixture consisted of retrovirus encoding a non-targeting control gRNA vector. For in vivo experiments, 5 × 10⁴ transduced P14 cells were intravenously transferred into Cas9-expressing, puromycin-resistant C57BL/6 recipient mice infected one day prior with either LCMV-Clone 13 or LCMV-Armstrong strain. A total of 25 LCMV-Clone 13 infected mice were used for five biological replicates, and 10 LCMV-Armstrong- infected mice were used for three biological replicates. Each biological replicate was labeled using hashtag antibodies (BioLegend, TotalSeq-C) to enable sample demultiplexing and statistical analysis. At ≥18 days post-infection, donor-derived P14 CD8^+^ T cells were sorted and pooled for Perturb-seq analysis. Preliminary tests indicated that T cells expressing gRNA *in vivo* exhibit a greater tendency for gRNA silencing over extended periods compared to *ex vivo* cultured cells, despite initial successful KOs. To mitigate gRNA barcode silencing, we harvested tissue between days 18 and 23. Sorted EGFP^+^ P14 CD8^+^ T cells were resuspended and diluted in 10% FBS RPMI at a concentration of 1 × 10^6^ cells per ml. Both the gene expression library and the CRISPR screening library were prepared using a Chromium Next GEM Single Cell 5′ kit with Feature Barcode technology for CRISPR Screening (10x Genomics). In brief, the single-cell suspensions were loaded onto the Chromium Controller according to their respective cell counts to generate 10,000 single-cell gel beads in emulsion per sample. Each sample was loaded into four separate channels. Chromium Next GEM Single Cell 5’ Kit v2, PN-1000263, Chromium 5’ Feature Barcode Kit, PN-1000541, 5’ CRISPR Kit, PN-1000451, Chromium Next GEM Chip K Single Cell Kit, PN-1000287, Dual Index Kit TT Set A, PN-1000215, Dual Index Kit TN Set A, PN- 1000250 (10x Genomics) in total were used for each reaction. The resulting libraries were quantified and quality checked using TapeStation (Agilent). Samples were diluted and loaded onto a NovaSeq (Illumina) using 100 cycle kit to obtain a minimum of 20,000 paired-end reads (26x91 bp) per cell for the gene expression library and 5,000 paired-end reads per cell for the CRISPR screening library, yielding an average of 42639, 36739, and 53413 reads aligned from cells from *in vivo* LCMV-Clone 13, *in vivo* LCMV-Armstrong infection, and *in vitro* donor respectively.

*Data analysis.* Alignments and count aggregation of gene expression and sgRNA reads were completed using Cell Ranger (v.7.0.1). Gene expression and sgRNA reads were aligned using the cell Ranger multi count command with default settings. Gene expression reads were aligned to the mouse genome (mm10 from ENSEMBL GRCm38 loaded from 10x Genomics). The median average of 4, 2, and 33 unique molecular identifiers (UMIs) were detected from cells from *in vivo* LCMV-Clone 13 and LCMV-Armstrong infection, and *in vitro* donor respectively. Droplets with sgRNA UMI passing of default Cell Ranger CRISPR analysis Protospacer UMI threshold were used in further analysis. The filtered feature matrices were imported into Seurat (v.4.3.0)^69^ to create assays for a Seurat object containing both gene expression and CRISPR guide capture matrices. Cells detected with sgRNAs targeting two or more genes were then removed to avoid interference from multi-sgRNA-transduced cells. Low-quality cells with <200 detected genes, >10% mitochondrial reads, and <5% ribosomal reads were discarded. A total of 17,257 cells (Clone 13) and 15,211 cells (Armstrong) were passed quality filtering and were used for downstream analysis. Count data was normalized by a global-scaling normalization method and linear transformed.^70^ Cluster-specific genes were identified using the FindAllMarkers function of Seurat. We utilized Nebulosa^71^ to recover signals from sparse features in single-cell data and made gRNA density plots with scCustomize^72^ based on kernel density estimation. In each biological replicate (Clone 13 n=5 and Armstrong =3), the percent cluster distribution of cells with each TF gRNA vector was calculated. Among two gRNA vectors per target TF, the gRNA vector with higher TEX_term_ reduction was shown in Fig. 3d and utilized for Perturb-seq in LCMV-Armstrong infection (Supplementary Table 5). Two-way ANOVA with Fisher’s LSD test was performed to determine statistical significance. DEG was calculated using the MAST model^73^, the results were then used as inputs for GSEA to evaluate the effect on selected pathways. Genes with p-value < 0.05 were considered as DEG.

UMAP plots were generated by calculating UMAP embeddings using Seurat and then plotting them as scatter plots using ggplot2. Kernel density calculations for each gRNA were performed on UMAP embeddings using the MASS package via the kde2d function. The kernel density results were plotted as a raster layer with ggplot2 over the UMAP scatter plots. Finally, density contour lines were added using ggplot2’s built-in 2D kernel density contour geom (geom_density_2d).

### ATAC-Seq library preparation and sequencing

ATAC-seq was performed as previously described^74^. Briefly, 5,000-50,000 viable cells were washed with cold PBS, collected by centrifugation, then lysed in resuspension buffer (RSB) (10 mM Tris-HCl, pH 7.4, 10 mM NaCl, 3 mM MgCl2) supplemented with 0.1% NP40, 0.1% Tween-20, and 0.01% digitonin. Samples were incubated on ice for 3 min, then washed out with 1 ml RSB containing 0.1% Tween-20. Nuclei were pelleted by centrifugation at 500g for 10 min at 4°C then resuspended in 50 ul transposition mix (25ul 2x TD buffer, 2.5 ul transposase (100 nM final), 16.5 ul PBS, 0.5 ul 1% digitonin, 0.5 ul 10% Tween-20, 5 ul H2O) and incubated at 37°C for 30 min in a thermomixer with 1000 RPM mixing. DNA was purified using a Qiagen MinElute PCR cleanup kit, then PCR amplified using indexed oligos. The optimal number of amplification cycles for each sample was determined by qPCR. Libraries were size selected using AmpureXP beads and sequenced using an Illumina NextSeq500 for 75bp paired-end reads.

### ATAC-Seq analysis

Paired-end 42-bp, or paired-end 75-bp reads were aligned to the M. musculus mm10 genome using BWA^75,76^with parameters “bwa mem -M -k 32”. ATAC-seq peaks were called using MACS2^77^ program using parameters “callpeaks -qvalue 5.0e-2 –shift -100 –extsize 200”. Differentially accessible regions were identified using DESeq2^78^ . Batch effect was removed using limma^79^. Heatmap visualization of ATAC-seq data was performed using pheatmap.

### Single-cell RNA sequencing metadata analysis

Analysis was primarily performed in R (v 3.6.1) using the package Seurat (v 3.1)^69,80^, with the package tidyverse (v 1.2.1)^81^ used to organize data and the package ggplot2 (v 3.2.1) used to generate figures. scRNA-seq data from GSE10898, GSE99254, GSE98638, GSE199565, and GSE181785 were filtered to keep cells with a low percentage of mitochondrial genes in the transcriptome (< 5%) and between 200 and 3000 unique genes to exclude poor quality reads and doublets. Cell cycle scores were regressed when scaling gene expression values and T cell receptor genes were regressed during the clustering process, which was performed with the Louvain algorithm within Seurat and visualized with UMAP. To quantify the gene expression patterns, we used Seurat’s module score feature to score each cluster based on its per cell expression of TFs.

To obtain Extended Data Fig. 5a, raw single-cell count data and cell annotation data was downloaded from NCBI GEO (GSE99254)^45^. Count data was normalized and transformed by derivation of the residuals from a regularized negative binomial regression model for each gene (SCT normalization method in Seurat^87^, v4.1.1), with 5000 variable features retained for downstream dimensionality reduction techniques. Integration of data was performed on the patient level with Canonical Correlation Analysis as the dimension reduction technique^82^. Principal component analysis and UMAP dimension reduction were performed, with the first 50 PCs utilized in UMAP generation. Cells were clustered utilizing the Louvain algorithm with multi-level refinement. The data was subset to CD8+ T cells, which were identified utilizing the labels provided by Guo et al^66^. Cell type labels were confirmed via 1) SingleR^83^ (v1.8.1) annotation utilizing the ImmGen^84^ database obtained via celda(1.10), 2) cluster marker identification, and 3) cell type annotation with the ProjecTILs T cell atlas^7^ (v2.2.1). After sub- setting to CD8^+^ T cells, cells were again normalized utilizing SCT normalization, with 3000 variable features retained for dimension reduction. Due to the low number of cells on the per-patient level, HArmstrongony^85^(v1.0) was utilized to integrate the data at the patient level, rather than Seurat. PCA and UMAP dimensionality reduction were performed as above.

### Statistical analyses

Statistical tests for flow cytometry data were performed using Graphpad Prism 10. *p-values* were calculated using either two-tailed unpaired Student’s t-tests, one-way ANOVA or two-way ANOVA as indicated in each figure. Linear regressions were performed using the ordinary least squares method in R (v 3.6.1). All data were presented as the mean ± s.e.m. The *P* values were represented as follows: \*\*\*\**P* < 0.0001, \*\*\**P* < 0.001, \*\**P* < 0.01, and \**P* < 0.05.

## Data availability

ATAC-sequencing data from this paper will be deposited in the GEO database (GSE279498). Taiji v2.0 output of this study (TF activity atlas, TF-TF interaction maps, and TF activity on genome browser view) will be available at our CD8^+^ T cell TF atlas portal (https://wangweilab.shinyapps.io/Tcellstates/) and interactive interface for TF atlas exploration (https://huggingface.co/spaces/taijichat/chat).

All other raw data are available from the corresponding author upon request.

## Code availability

All scripts and the Taiji v2.0 package are uploaded to GitHub (https://github.com/Wang-lab-UCSD/Taiji2).

---

**Extended Data Fig. 1.** | Parallel differentiation of T_RM_ and TEX_term_ and their transactional and epigenetic similarity. a, UMAP of scRNA-seq data of T cells from blood, tumor, and adjacent normal tissues of CRC^86^, NSCLC^45^, and HCC^61^ patients. Unbiased clustering identified multiple T cell states consistent with those observed murine LCMV infection and tumors. **b,** T_RM_ marker genes show higher expression in TEX_term_ cluster from Pan-cancer scRNA-seq in **a**. **c,** Both T_RM_ and TEX_term_ clusters upregulate exhaustion-^8^ and T ^35^-associated gene signatures. **d,** Pearson correlation matrix of batch- corrected ATAC-seq datasets^3,9,33,34^. Color and size are both proportional to correlation strength. **e,** A total of 121 experiments across multiple data sets^3,9,18,23,32–36^ were utilized to generate an epigenetic and transcriptional atlas of CD8^+^ T cells under chronic and acute antigen exposure.

**Extended Data Fig. 2.** | Cataloging key TFs across CD8^+^ T cell states. a, Logic flow of the unbiased PageRank comparison to classify single-state/multi-state TFs. **b,** Number of TFs catalogued in each cell state. **c,** Venn Diagrams showing overlap of TFs with the T_RM_ cell state. **d,** TF activity score (normalized PageRank) of previously reported TEX_term_-preventing TFs (TFAP4, JUN, BATF3, and BATF), newly identified TEX_term_ single-state TFs (JDP2, ZFP324, ZBTB49, ZFP143, ZSCAN20), and NFATC1, a known TEX_term_-associated TF.

**Extended Data Fig. 3.** | TF wave analysis. a, Schematic of the analysis pipeline. **b,** Selection of algorithms and parameters for TF wave analysis. The Pearson correlation was chosen for the distance metric, with k=7 chosen as the optimal cluster number. **c,** Seven TF waves. Circles represent specific cell states. Red color indicates normalized PageRank scores. **d,** List of TF members in each wave. **e**, Heatmap of biological pathways enriched in each TF wave. Red-blue color scale indicates the p-value.

**Extended Data Fig. 4.** | TF–TF association network in TEX_term_ and T_RM_ cell states. a–c, TF–TF associations of (**a**) TEX_term_ single-state TFs (JDP2, ZFP324, and IRF8); (**b**) T_RM_ single-state TFs (FOSB, SNAI1, and KLF6); (**c**) multi-state TFs shared by TEX_term_ and T_RM_ (PRDM1, FLI1, and GFI1); and (**d**) previously reported TFs whose OE prevent TEXterm–JUN, TFAP4, and BATF. Line color indicates state specificity: T_RM_ (green) or TEX_term_ (brown) state. Line thickness represents interaction intensity. Line color indicates state specificity: T_RM_ (green) or TEX_term_ (brown) state. Line thickness represents interaction intensity.

**Extended Data Fig. 5.** | TF network analysis reveals proteasome pathway enrichment in TEX_term_ state with diminished tumor control function. a, Pseudotime analysis of CD8^+^ TIL scRNA-seq data from NSCLC patients (n = 14), showing a positive correlation between proteasome gene scores (KEGG: M10680) and T cell exhaustion. b, Gene set enrichment analysis (GO:0043161, Proteasome-mediated ubiquitin-dependent protein catabolic process) of RNA-seq from CD8^+^ splenocytes (black) and TILs (purple), n = 3 each. c, Tumor growth of Rag2^-/-^ mice bearing MCA-205 sarcomas infused with proteasome^high^ or proteasome^low^ CD8^+^ TILs isolated from C57BL/6 mice bearing MCA-205 tumors. n = 5 per group. Two-way ANOVA Tukey’s multiple comparison test were performed. *\*\*\*\*P*<0.0001, *\*\*\*P*<0.001, *\*\*P*<0.01, *\*P*<0.05.

**Extended Data Fig. 6.** | *in vivo* Perturb-seq with dual-guide RNA in LCMV chronic infection and individual validation. a, Retroviral vector design for dual-gRNA delivery. **b,** Feature plot of differentiation markers (Pan-exhaustion: red, TEX_prog_: pink, TEX_eff_: blue, Cell cycle: purple, TEX_term_: brown). **c,** Heatmap of marker gene expression across cell state clusters identified by Seurat’s Find Markers() function. **d,** Frequency of PD1^+^ SLAMF6^−^ CD101^+^ cells (**d**). Frequency of CD38^+^ CD39^+^ double-positive cells (**e**). Statistical analysis: Ordinary one-way ANOVA with Dunnett’s multiple comparisons test versus gScramble (n ≥ 5 from ≥2 biological replicates). Data are presented as mean ± s.e.m. *\*\*\*\*P<* 0.0001; *\*\*\*P <* 0.001; *\*\*P <* 0.01; *\*P <* 0.05.

**Extended Data Fig. 7.** | Single-cell transcriptomic profiling of CD8^+^ T cells from Perturb-seq in acute LCMV infection and validation of the selectivity of T_RM_ single-state TF, *Klf6* overexpression. a, Feature plots of differentiation marker genes (T_CM_: yellow, T_EM_: blue, T_RM_-*Itgae*^low^: dark blue, T_RM_: green). b, Heatmap of differentially expressed genes between T_RM_-*Itgae*^low^ and T_RM_ clusters. c, Volcano plots of differentially expressed genes in Cas9^+^ P14 CD8^+^ T cells expressing *gEtv5*, *gArid3a*, g*Hic1*, or *gGfi1*. d, T_RM_ single-state TF, *Klf6* overexpression does not accelerate T cell exhaustion. Experimental setup: *Klf6-*RV or control-RV transduced P14 CD8^+^ T cells co-transferred into mice infected with chronic LCMV-Clone 13, Quantification of the frequency of PD1^+^, TEX_prog_ (PD1^+^ SLAMF6^+^ CX3CR1^-^), TEX_eff_ (PD1^+^ SLAMF6^-^ CX3CR1^+^) and TEX_term_ (PD1^+^ SLAMF6^-^CX3CR1^-^) populations. Paired t-tests (n ≥ 6 from ≥2 biological replicates). Data are presented as mean ± s.e.m. *\*\*\*\*P <*0.0001, *\*\*\*P <* 0.001, *\*\*P <* 0.01, *\*P <* 0.05.

**Extended Data Fig. 8.** | Depletion of TEX_term_-single state TF, *Zscan20* and TEX_term_ and T_RM_ multi- state TF, *Hic1* reduces T cell exhaustion. a, Co-transfer of Cas9^+^ P14 CD8^+^ T cells transduced with RV-g*Zscan20* or RV-g*Hic1*, mixed with gRNA control RV transduced cells into B16-GP33 tumor- bearing mice. b, Representative flow plots of PD1^+^ P14 CD8^+^ T cells stained for SLAMF6 and CX3CR1. c-e, Quantification of (c) TEX_prog_ (PD1^+^ SLAMF6^+^ TIM3⁻), (d) TEX_term_ (PD1^+^ SLAMF6⁻ CX3CR1⁻), and (e) non-progenitor, exhausted (PD1^+^ SLAMF6⁻ TIM3^+^) popultation. Paired t-tests (n ≥ 6 from ≥2 biological replicates). Data are presented as mean ± s.e.m. ****P < 0.0001, ***P < 0.001, **P < 0.01, *P < 0.05.

**Extended Data Fig. 9.** | Conservation of single- and multi-state TFs in human pan-cancer TEX_term_ and T_RM_ cell states. a, Human pan-cancer datasets utilized in this study^49–56^. **b,** Cluster-specific marker mRNA expression across integrated pan-cancer single-cell multi-omics datasets. **c,** TF activity analysis using Taiji on matched matched scRNA-seq and scATAC-seq datasets from various human cancers, including ccRCC, GBM, BCC, HNSCC, HCC, and RCC. CD8^+^ T cell states were annotated using canonical marker gene expression. PageRank scores derived from Taiji were log-transformed, averaged per state, and z-score normalized. Results were visualized with a focus on TF activity T_RM_ and TEX_term_ cell states. **d,** mRNA expression of TFs were compared across human CD8^+^ T cell states. The dataset encompassed T cells from 15 tumor types. Cell type annotations provided by the original study were retained and mapped to the following broad categories of clusters: T_RM_, TEX_term_, T_EM_, Memory, Naive, and cell cycle. Seurat’s AverageExpression function was used, followed by z-score normalization. Data visualization emphasized comparisons between T_RM_ and TEX_term_ cell states. Cell-state-selectivity conserved TFs in humans and mice are highlighted in bold (**c, d**).

**Extended Data Fig. 10.** | CRISPR validation of TEX_term_ single-state TF KO in human PBMCs and antitumor activity of TF-deficient T cells in mice. Indel frequencies, representative Sanger sequencing traces, and indel distributions for (a) *ZSCAN20* and (b) *JDP2* KOs. c, Survival of B16-GP33 bearing mice receiving *Zscan20* KO or control P14 CD8^+^ T cells, followed by anti-PD1 or isotype IgG2a treatment. d, Survival of tumor-bearing mice with *Jdp2* KO P14 CD8^+^ T cell transfer.

**Supplementary Table 1.** | Composition of multi-omic atlas

**Supplementary Table 2.** | TF PageRank scores of multi-state and single-state TFs

**Supplementary Table 3.** | TF PageRank scores of T_RM_- and TEX_term_ single-state TFs and multi-state TFs

**Supplementary Table 4.** | Catalog of cell-state important TFs

**Supplementary Table 5.** | Catalog of TF communities in T_RM_ and TEX_term_

**Supplementary Table 6.** | gRNA sequences for TF gRNA library and RNP gRNA

**Supplementary Table 7.** | Perturb seq cluster statistics (Fig. 3 and Fig. 4)

**Supplementary Table 8.** | Gene signatures

## Supplementary Methods

TaijiChat: An Integrated Conversational Interface for Multi-Omics Data Exploration.

## References

1. Chung, H. K., McDonald, B. & Kaech, S. M. The architectural design of CD8+ T cell responses in acute and chronic infection: Parallel structures with divergent fates. J. Exp. Med. 218, (2021).

2. Kasmani, M. Y. et al. Clonal lineage tracing reveals mechanisms skewing CD8+ T cell fate decisions in chronic infection. J. Exp. Med. 220, (2023).

3. Beltra, J.-C. et al. Developmental Relationships of Four Exhausted CD8 T Cell Subsets Reveals Underlying Transcriptional and Epigenetic Landscape Control Mechanisms. Immunity 52, 825– 841.e8 (2020).

4. Giles, J. R. et al. Shared and distinct biological circuits in effector, memory and exhausted CD8 T cells revealed by temporal single-cell transcriptomics and epigenetics. Nature Immunology vol. 23 1600–1613 Preprint at 10.1038/s41590-022-01338-4 (2022).

5. Pauken, K. E. et al. Epigenetic stability of exhausted T cells limits durability of reinvigoration by PD-1 blockade. Science 354, 1160–1165 (2016).

6. Sen, D. R. et al. The epigenetic landscape of T cell exhaustion. Science vol. 354 1165–1169 Preprint at 10.1126/science.aae0491 (2016).

7. Wherry, E. J., Blattman, J. N., Murali-Krishna, K., van der Most, R. & Ahmed, R. Viral persistence alters CD8 T-cell immunodominance and tissue distribution and results in distinct stages of functional impairment. J. Virol. 77, 4911–4927 (2003).

8. Wherry, E. J. et al. Molecular signature of CD8+ T cell exhaustion during chronic viral infection. Immunity 27, 670–684 (2007).

9. Milner, J. J. et al. Runx3 programs CD8 T cell residency in non-lymphoid tissues and tumours. Nature 552, 253–257 (2017).

10. Ganesan, A.-P. et al. Tissue-resident memory features are linked to the magnitude of cytotoxic T cell responses in human lung cancer. Nat. Immunol. 18, 940–950 (2017).

11. Djenidi, F. et al. CD8 CD103 Tumor–Infiltrating Lymphocytes Are Tumor-Specific Tissue- Resident Memory T Cells and a Prognostic Factor for Survival in Lung Cancer Patients. The Journal of Immunology vol. 194 3475–3486 Preprint at 10.4049/jimmunol.1402711 (2015).

12. Corgnac, S., Boutet, M., Kfoury, M., Naltet, C. & Mami-Chouaib, F. The Emerging Role of CD8+ Tissue Resident Memory T (TRM) Cells in Antitumor Immunity: A Unique Functional Contribution of the CD103 Integrin. Front. Immunol. 9, 1904 (2018).

13. Komdeur, F. L. et al. CD103+ tumor-infiltrating lymphocytes are tumor-reactive intraepithelial CD8+ T cells associated with prognostic benefit and therapy response in cervical cancer. Oncoimmunology 6, e1338230 (2017).

14. Kaech, S. M. & Cui, W. Transcriptional control of effector and memory CD8+ T cell differentiation. Nat. Rev. Immunol. 12, 749–761 (2012).

15. Sen, D. R. et al. The epigenetic landscape of T cell exhaustion. Science vol. 354 1165–1169 Preprint at 10.1126/science.aae0491 (2016).

16. Im, S. J. et al. Defining CD8+ T cells that provide the proliferative burst after PD-1 therapy. Nature 537, 417–421 (2016).

17. Miller, B. C. et al. Subsets of exhausted CD8 T cells differentially mediate tumor control and respond to checkpoint blockade. Nat. Immunol. 20, 326–336 (2019).

18. Hudson, W. H. et al. Proliferating Transitory T Cells with an Effector-like Transcriptional Signature Emerge from PD-1 Stem-like CD8 T Cells during Chronic Infection. Immunity 51, 1043–1058.e4 (2019).

19. Liu, F. et al. CTLA-4 correlates with immune and clinical characteristics of glioma. Cancer Cell Int. 20, 7 (2020).

20. Zhang, M. et al. Prognostic Values of CD38+CD101+PD1+CD8+ T Cells in Pancreatic Cancer. Immunol. Invest. (2019) doi:10.1080/08820139.2019.1566356.

21. Shin, H. et al. A role for the transcriptional repressor Blimp-1 in CD8(+) T cell exhaustion during chronic viral infection. Immunity 31, 309–320 (2009).

22. Behr, F. M. et al. Blimp-1 Rather Than Hobit Drives the Formation of Tissue-Resident Memory CD8 T Cells in the Lungs. Front. Immunol. 10, 400 (2019).

23. Milner, J. J. et al. Heterogenous Populations of Tissue-Resident CD8+ T Cells Are Generated in Response to Infection and Malignancy. Immunity 52, 808–824.e7 (2020).

24. Li, C. et al. The Transcription Factor Bhlhe40 Programs Mitochondrial Regulation of Resident CD8 T Cell Fitness and Functionality. Immunity 51, 491–507.e7 (2019).

25. Wu, J. E. et al. In vitro modeling of CD8 T cell exhaustion enables CRISPR screening to reveal a role for BHLHE40. Sci Immunol 8, eade3369 (2023).

26. Chen, J. et al. NR4A transcription factors limit CAR T cell function in solid tumours. Nature 567, 530–534 (2019).

27. Mackay, L. K. et al. The developmental pathway for CD103(+)CD8+ tissue-resident memory T cells of skin. Nat. Immunol. 14, 1294–1301 (2013).

28. Schacht, T., Oswald, M., Eils, R., Eichmüller, S. B. & König, R. Estimating the activity of transcription factors by the effect on their target genes. Bioinformatics 30, i401–7 (2014).

29. Zhang, K., Wang, M., Zhao, Y. & Wang, W. Taiji: System-level identification of key transcription factors reveals transcriptional waves in mouse embryonic development. Sci Adv 5, eaav3262 (2019).

30. Wang, J., Liu, C., Chen, Y. & Wang, W. Taiji-reprogram: a framework to uncover cell-type specific regulators and predict cellular reprogramming cocktails. NAR Genom Bioinform 3, lqab100 (2021).

31. Liu, C. et al. Systems-level identification of key transcription factors in immune cell specification. PLoS Comput. Biol. 18, e1010116 (2022).

32. Milner, J. J. et al. Delineation of a molecularly distinct terminally differentiated memory CD8 T cell population. Proc. Natl. Acad. Sci. U. S. A. 117, 25667–25678 (2020).

33. Guan, T. et al. ZEB1, ZEB2, and the miR-200 family form a counterregulatory network to regulate CD8 T cell fates. J Exp Med 215, 1153–1168 (2018).

34. Scott-Browne, J. P. et al. Dynamic Changes in Chromatin Accessibility Occur in CD8 T Cells Responding to Viral Infection. Immunity 45, 1327–1340 (2016).

35. Mackay, L. K. et al. Hobit and Blimp1 instruct a universal transcriptional program of tissue residency in lymphocytes. Science 352, 459–463 (2016).

36. Renkema, K. R. et al. KLRG1 Memory CD8 T Cells Combine Properties of Short-Lived Effectors and Long-Lived Memory. J Immunol 205, 1059–1069 (2020).

37. Intlekofer, A. M. et al. Effector and memory CD8+ T cell fate coupled by T-bet and eomesodermin. Nat Immunol 6, 1236–1244 (2005).

38. Reiser, J., Sadashivaiah, K., Furusawa, A., Banerjee, A. & Singh, N. Eomesodermin driven IL- 10 production in effector CD8 T cells promotes a memory phenotype. Cell Immunol 335, 93–102 (2019).

39. Crowl, J. T. et al. Tissue-resident memory CD8 T cells possess unique transcriptional, epigenetic and functional adaptations to different tissue environments. Nature Immunology vol. 23 1121–1131 Preprint at 10.1038/s41590-022-01229-8 (2022).

40. Chandele, A. et al. Formation of IL-7Ralphahigh and IL-7Ralphalow CD8 T cells during infection is regulated by the opposing functions of GABPalpha and Gfi-1. J. Immunol. 180, 5309–5319 (2008).

41. Schmidt, R. et al. CRISPR activation and interference screens decode stimulation responses in primary human T cells. Science 375, eabj4008 (2022).

42. Shifrut, E. et al. Genome-wide CRISPR Screens in Primary Human T Cells Reveal Key Regulators of Immune Function. Cell 175, 1958–1971.e15 (2018).

43. McCutcheon, S. R. et al. Transcriptional and epigenetic regulators of human CD8+ T cell function identified through orthogonal CRISPR screens. Nat. Genet. 55, 2211–2223 (2023).

44. Seo, H. et al. BATF and IRF4 cooperate to counter exhaustion in tumor-infiltrating CAR T cells. Nat. Immunol. 22, 983–995 (2021).

45. Guo, X. et al. Global characterization of T cells in non-small-cell lung cancer by single-cell sequencing. Nat Med 24, 978–985 (2018).

46. Riesenberg, B. P. et al. Stress-Mediated Attenuation of Translation Undermines T-cell Activity in Cancer. Cancer Res. 82, 4386–4399 (2022).

47. Daniel, B. et al. Divergent clonal differentiation trajectories of T cell exhaustion. Nat. Immunol. 23, 1614–1627 (2022).

48. Lin, Y. H. et al. Small intestine and colon tissue-resident memory CD8+ T cells exhibit molecular heterogeneity and differential dependence on Eomes. Immunity 56, 207–223.e8 (2023).

49. Long, Z. et al. Single-cell multiomics analysis reveals regulatory programs in clear cell renal cell carcinoma. Cell Discov 8, 68 (2022).

50. Terekhanova, N. V. et al. Epigenetic regulation during cancer transitions across 11 tumour types. Nature 623, 432–441 (2023).

51. Riegel, D. et al. Integrated single-cell profiling dissects cell-state-specific enhancer landscapes of human tumor-infiltrating CD8 T cells. Mol Cell 83, 622–636.e10 (2023).

52. Yost, K. E. et al. Clonal replacement of tumor-specific T cells following PD-1 blockade. Nat Med 25, 1251–1259 (2019).

53. Ma, L. et al. Tumor Cell Biodiversity Drives Microenvironmental Reprogramming in Liver Cancer. Cancer Cell 36, 418–430.e6 (2019).

54. Young, M. D. et al. Single-cell transcriptomes from human kidneys reveal the cellular identity of renal tumors. Science 361, 594–599 (2018).

55. Cillo, A. R. et al. Immune Landscape of Viral- and Carcinogen-Driven Head and Neck Cancer. Immunity 52, 183–199.e9 (2020).

56. Zheng, L. et al. Pan-cancer single-cell landscape of tumor-infiltrating T cells. Science 374, abe6474 (2021).

57. Chen, Z. et al. In vivo CD8+ T cell CRISPR screening reveals control by Fli1 in infection and cancer. Cell 184, 1262–1280.e22 (2021).

58. Zhou, P. et al. Single-cell CRISPR screens in vivo map T cell fate regulomes in cancer. Nature 624, 154–163 (2023).

59. Van Der Byl, W. et al. The CD8+ T cell tolerance checkpoint triggers a distinct differentiation state defined by protein translation defects. Immunity (2024) doi:10.1016/j.immuni.2024.04.026.

60. Chang, J. T. et al. Asymmetric proteasome segregation as a mechanism for unequal partitioning of the transcription factor T-bet during T lymphocyte division. Immunity 34, 492–504 (2011).

## Methods references

61. Zheng, C. et al. Landscape of Infiltrating T Cells in Liver Cancer Revealed by Single-Cell Sequencing. Cell 169, 1342–1356.e16 (2017).

62. Weirauch, M. T. et al. Determination and Inference of Eukaryotic Transcription Factor Sequence Specificity. Cell vol. 158 1431–1443 Preprint at 10.1016/j.cell.2014.08.009 (2014).

63. Zhao, T., Liu, H., Roeder, K., Lafferty, J. & Wasserman, L. The huge Package for High-dimensional Undirected Graph Estimation in R. (2020) doi:10.48550/arXiv.2006.14781.

64. Traag, V. A., Waltman, L. & van Eck, N. J. From Louvain to Leiden: guaranteeing well-connected communities. Sci. Rep. 9, 5233 (2019).

65. Schönfeld, M. & Pfeffer, J. Fruchterman/Reingold (1991): Graph Drawing by Force-Directed Placement. Schlüsselwerke der Netzwerkforschung 217–220 Preprint at 10.1007/978-3-658-21742-6_49 (2019).

66. Bailis, W. et al. Author Correction: Distinct modes of mitochondrial metabolism uncouple T cell differentiation and function. Nature 573, E2 (2019).

67. Cong, L. et al. Multiplex genome engineering using CRISPR/Cas systems. Science 339, 819–823 (2013).

68. Wang, B. et al. Integrative analysis of pooled CRISPR genetic screens using MAGeCKFlute. Nat. Protoc. 14, 756–780 (2019).

69. Stuart, T. et al. Comprehensive Integration of Single-Cell Data. Cell vol. 177 1888–1902.e21 Preprint at 10.1016/j.cell.2019.05.031 (2019).

70. Hafemeister, C. & Satija, R. Normalization and variance stabilization of single-cell RNA-seq data using regularized negative binomial regression. Genome Biol. 20, 296 (2019).

71. Alquicira-Hernandez, J. & Powell, J. E. Nebulosa recovers single-cell gene expression signals by kernel density estimation. Bioinformatics 37, 2485–2487 (2021).

72. Marsh, S. E. scCustomize: custom visualizations & functions for streamlined analyses of single cell sequencing. Preprint at 10.5281/zenodo.

73. Finak, G. et al. MAST: a flexible statistical framework for assessing transcriptional changes and characterizing heterogeneity in single-cell RNA sequencing data. Genome Biol. 16, 278 (2015).

74. McDonald, B. et al. Canonical BAF complex activity shapes the enhancer landscape that licenses CD8 T cell effector and memory fates. Immunity 56, 1303–1319.e5 (2023).

75. Li, H. & Durbin, R. Fast and accurate short read alignment with Burrows-Wheeler transform. Bioinformatics 25, (2009).

76. Li, H. & Durbin, R. Fast and accurate long-read alignment with Burrows-Wheeler transform. Bioinformatics 26, (2010).

77. Zhang, Y. et al. Model-based Analysis of ChIP-Seq (MACS). Genome Biol. 9, 1–9 (2008).

78. Love, M. I., Huber, W. & Anders, S. Moderated estimation of fold change and dispersion for RNA- seq data with DESeq2. Genome Biol. 15, 1–21 (2014).

79. Ritchie, M. E. et al. limma powers differential expression analyses for RNA-sequencing and microarray studies. Nucleic Acids Res. 43, e47 (2015).

80. Butler, A., Hoffman, P., Smibert, P., Papalexi, E. & Satija, R. Integrating single-cell transcriptomic data across different conditions, technologies, and species. Nat. Biotechnol. 36, 411–420 (2018).

81. Wickham, H. et al. Welcome to the Tidyverse. Journal of Open Source Software vol. 4 1686 Preprint at 10.21105/joss.01686 (2019).

82. Hao, Y. et al. Integrated analysis of multimodal single-cell data. Cell 184, 3573–3587.e29 (2021).

83. Aran, D. et al. Reference-based analysis of lung single-cell sequencing reveals a transitional profibrotic macrophage. Nat. Immunol. 20, 163–172 (2019).

84. Heng, T. S. P., Painter, M. W. & Immunological Genome Project Consortium. The Immunological Genome Project: networks of gene expression in immune cells. Nat. Immunol. 9, 1091–1094 (2008).

85. Korsunsky, I. et al. Fast, sensitive and accurate integration of single-cell data with Harmony. Nat. Methods 16, 1289–1296 (2019).

86. Zhang, L. et al. Single-Cell Analyses Inform Mechanisms of Myeloid-Targeted Therapies in Colon Cancer. Cell 181, 442–459.e29 (2020).

