## Extended Data Figures for "Atlas-Guided Discovery of Transcription Factors for T Cell Programming"

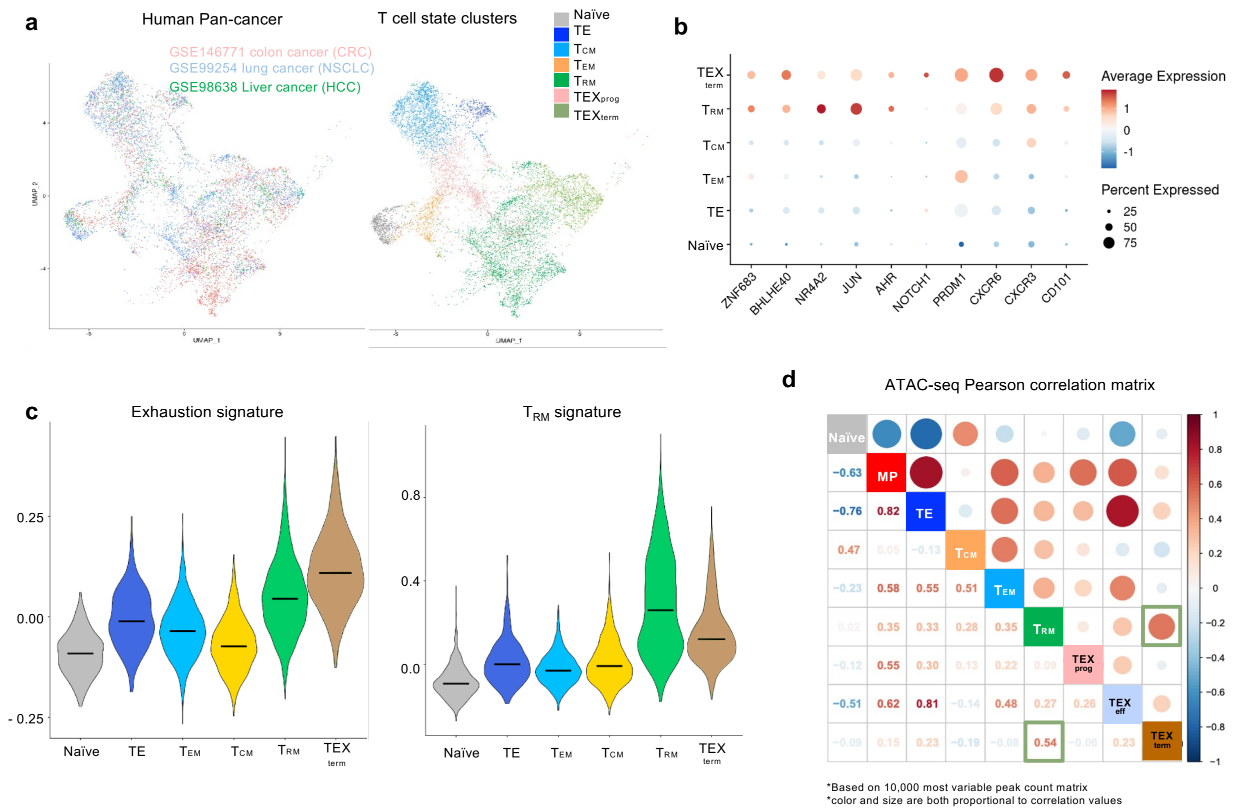


**e** 10 datasets | 9 subsets | 121 experiments | 52 ATAC-seq | 69 RNA-seq

| Author (Year) Accession | Data type | Infection | Naive | MP (MPEC) | TE (SLEC) | $T_{CM}$ | $T_{EM}$ | $T_{RM}$ | $TEX_{prog}$ | $TEX_{eff}$ | $TEX_{term}$ |
|---|---|---|---|---|---|---|---|---|---|---|---|
| Milner (2017) GSE107281 | RNA, ATAC | LCMV Arm | | | | | | Arm d7 CD8+ P14 CD69+ CD103+ (Small Intestine) | | | |
| Milner (2020.Apr) GSE147502 | RNA | LCMV Arm | | | | | | Arm d55 CD8+ P14 (Small Intestine) | | | |
| Milner (2020.Aug) GSE157072 | RNA | LCMV Arm | d0 CD8+ P14 | Arm d7 CD8+ P14 IL7R+ KLRG1- | Arm d7 CD8+ P14 IL7R- KLRG1+ | Arm d55 CD8+ P14 IL7R+ CD62L+ | Arm d55 CD8+ P14 IL7R+ CD62L- | | | | |
| Guan (2018) GSE111902 | RNA, ATAC | LCMV Arm | d0 CD8+ P14 | Arm d8 CD8+ P14 IL7R+ KLRG1- | Arm d8 CD8+ P14 IL7R- KLRG1+ | | | | | | |
| Scott-Browne (2016) GSE88987 | RNA, ATAC | LCMV Arm | d0 CD8+ P14 | Arm d8 CD8+ P14 IL7R+ KLRG1- (ATAC) | Arm d8 CD8+ P14 IL7R- KLRG1+ (ATAC) | | | | | | |
| Mackay (2016) GSE70813 | RNA | LCMV Arm | d0 CD8+ P14 | | | Arm d40+ CD8+ P14 CD62L+ CD69- | Arm d40+ CD8+ P14 CD62L- CD69- | Arm CD8+ P14 d40+ CD62L- CD69+ (Liver) | | | |
| Renkema (2020) GSE152841 | RNA | LCMV Arm | d0 CD8+ CD44- | | | Arm d90 CD8+ CD44+ CD62L+ KLRG1- CD27+ | Arm d90 CD8+ CD44+ CD62L- KLRG1- CD27+ | | | | |
| Hudson (2019) PRJNA497086 | RNA | LCMV CL13 | d0 | | | | | | CL13 d45+ CD8+ PD1+ CD101-Tim3- | CL13 d45+ CD8+ PD1+ CD101-Tim3+ | CL13 d45+ CD8+ PD1+ CD101+Tim3+ |
| Beltra (2020) GSE149879 | RNA, ATAC | LCMV CL13 | | | | | | | CL13 d30 CD8+ PD1+ SLAMF6+ | CL13 d30 CD8+ PD1+ SLAMF6- CD69- | CL13 d30 CD8+ PD1+ SLAMF6- CD69+ |
| This study GSE279498 | ATAC | LCMV CL13 | d0 CD8+ CD44- | Arm d7 CD8+ gp33+ gp276+ IL7R+ KLRG1- | Arm d7 CD8+ gp33+ gp276+ IL7R- KLRG1+ | Arm d30 CD8+ gp33+ gp276+ CD62L+ | Arm d30 CD8+ gp33+ gp276+ CD62L- | Arm CD8+ gp33+ gp276+ CD62L- (Liver, Small Intestine) | CL13 d45+ CD8+ gp33+ gp276+ PD1+ SLAMF6+ Tim3- | CL13 d45+ CD8+ gp33+ gp276+ PD1+ SLAMF6- Tim3+ CD101- | CL13 d45+ CD8+ gp33+ gp276+ PD1+ SLAMF6- Tim3+ CD101+ |

**Extended Data Fig. 1 | Parallel differentiation of $T_{RM}$ and $TEX_{term}$ and their transactional and epigenetic similarity. a,** UMAP of scRNA-seq data of T cells from blood, tumor, and adjacent normal tissues of CRC[99], NSCLC[47], and HCC[74] patients. Unbiased clustering identified multiple T cell states consistent with those observed murine LCMV infection and tumors. **b,** $T_{RM}$ marker genes show higher expression in $TEX_{term}$ cluster from Pan-cancer scRNA-seq in **a**. **c,** Both $T_{RM}$ and $TEX_{term}$ clusters upregulate exhaustion-[17] and $T_{RM}$[39]-associated gene signatures. **d,** Pearson correlation matrix of batch-corrected ATAC-seq datasets[3,9,37,38]. Color and size are both proportional to correlation strength. **e,** Composition of multi-omic atlas. A total of 121 experiments across multiple data sets[3,9,20,27,36–40] were utilized to generate an epigenetic and transcriptional atlas of CD8$^+$ T cells under chronic and acute antigen exposure.

a

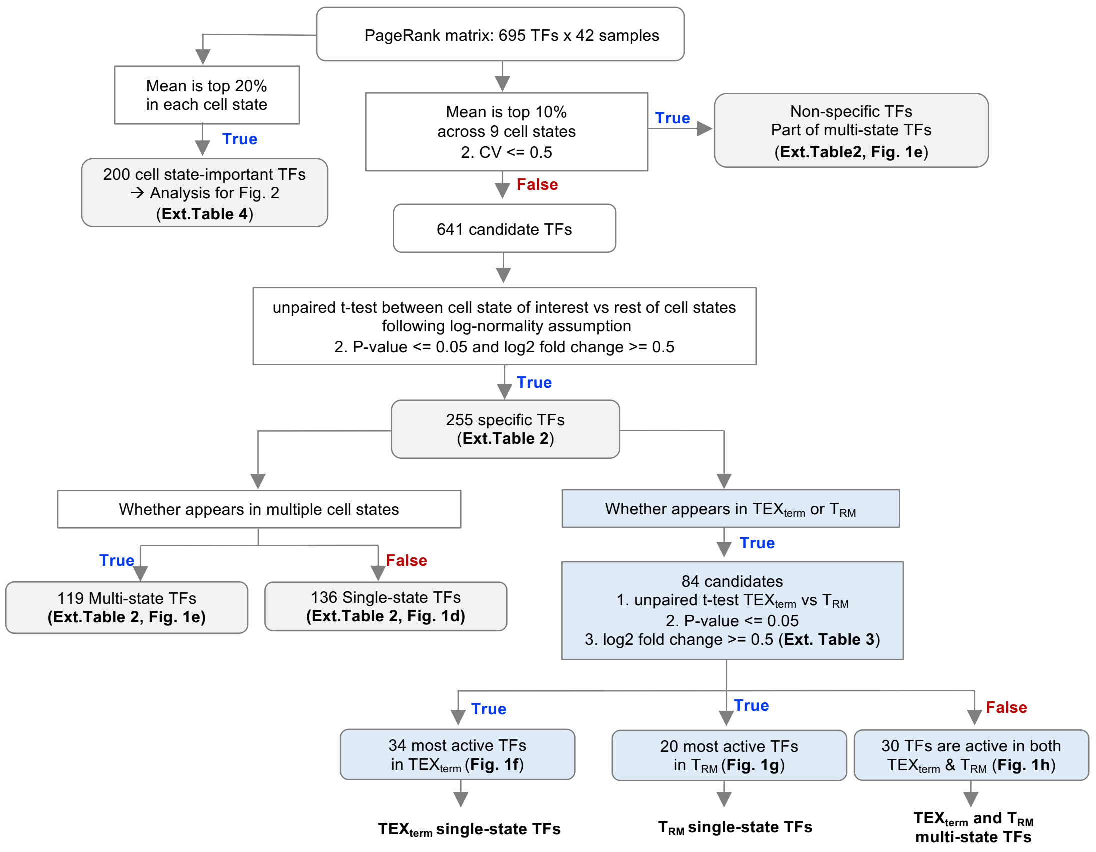


b

| | # of TFs active in each cell state | # of state-specific TFs | New, specific TFs examples |
|---|---|---|---|
| Non-specific TFs | 54 | NA | |
| Naive | 31 | 14 | *Hoxa7* |
| MP | 57 | 18 | *Hoxb4* |
| TE | 57 | 15 | *Amtl* |
| $T_{CM}$ | 28 | 15 | *Cux1* |
| $T_{EM}$ | 25 | 13 | *Meis3, Zfp398* |
| $T_{RM}$ | 43 | 17 | *Snai1, Klf6* |
| $TEX_{prog}$ | 63 | 19 | *Hey1* |
| $TEX_{eff}$ | 62 | 13 | *Sox8* |
| $TEX_{term}$ | 51 | 12 | *Zscan20, Zfp143, Zbtb49* |

c

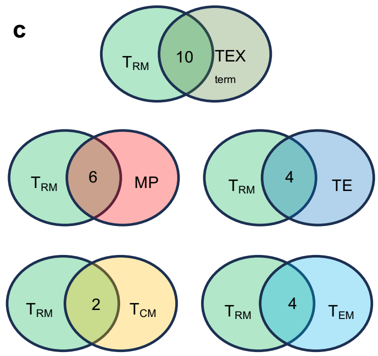


d

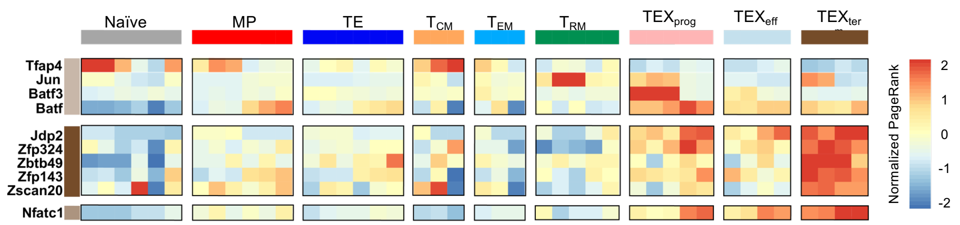


**Extended Data Fig. 2 | Cataloging key TFs across CD8$^+$ T cell states. a,** Logic flow of the unbiased PageRank comparison to classify single-state/multi-state TFs. **b,** Number of TFs catalogued in each cell state. **c,** Venn Diagrams showing overlap of TFs with the $T_{RM}$ cell state. **d,** TF activity score (normalized PageRank) of previously reported $TEX_{term}$-preventing TFs (TFAP4, JUN, BATF3, and BATF), newly identified $TEX_{term}$ single-state TFs (JDP2, ZFP324, ZBTB49, ZFP143, ZSCAN20), and NFATC1, a known $TEX_{term}$-associated TF.

**a**

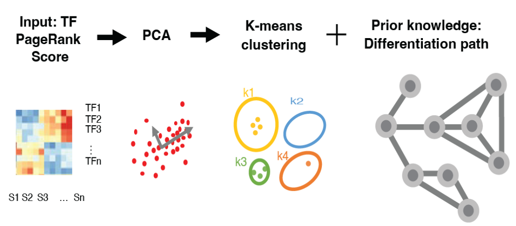


**b**

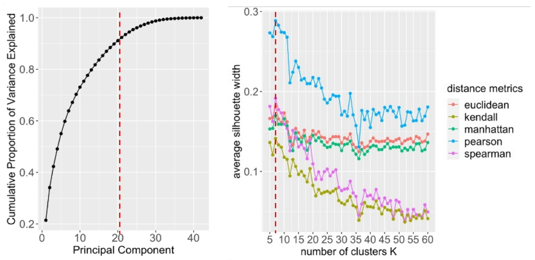


**c**

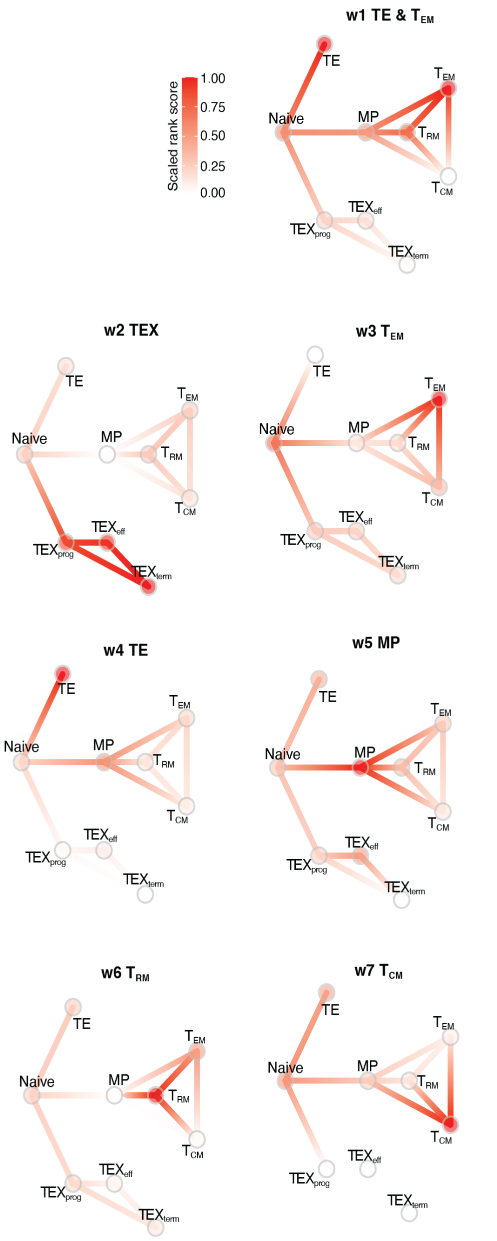


**d**

| Wave | TFs (Top 30) |
|---|---|
| **w1:** TE & $T_{EM}$ | *Sall4 Insm1 Ascl1 Sox17 Nr2f2 Nhlh1 Tfap2c Klf15 Gata4 Tfap2b Tal2 Gata6 Atf7 Esr2 Neurog2 Foxi3 Six1 Ehf Elf5 Sox3 Zfp454 Nhlh2 Foxa3 Prdm14 Gli3 Sp8 Tcfl5 Six4 Foxp2* |
| **w2:** TEX | *Vax2 Jdp2 Stat3 Irf8 Nfatc1 Irf9 Irf7 Klf10 Prdm16 Klf9 E2f6 Zfp324 Eomes Tfdp1 Stat2 Dmrt2 Stat1 Zfp41 Irf1 E2f4 Zfp523 Batf Zfp143 Grhl1 Stat5b Cebpb Spic Zfp128 Tfeb Osr2 Pax9* |
| **w3:** $T_{EM}$ | *Sp140 Atf6 Lcor Arntl E2f2 Naif1 Zfp69 Vezf1 Tbx21 Zfp580 Lbx2 Zzz3 Mybl1 Zfp189 Zfp212 Zfp260 Creb3l1 Lcorl Sp110 Nr6a1 Fli1 Ets1 Zkscan1 Hoxb1 Zfp932 Gabpa Hmg20b Mlxipl* |
| **w4:** TE | *Nr1d1 Tfap2a Foxo4 Zfp652 Nr1h2 Gli1 Zbtb48 Glis2 Smad3 E2f5 Phf21a Clock Hmbox1 Tead2 Bcl6b Bach2 Nr1i3 Zfp768 Rarb Taf1 Nr2c2 Zfp296 Zscan10 Nr1i2 Grhl2 Cux2 Tgif2* |
| **w5:** MP | *Lef1 Hoxa7 Sox4 Nfix Gata2 Esr1 Tcf7 Foxj3 Lyl1 Pbx1 Cebpa Tcf12 Foxj1 Hand2 Cebpe Gata1 Cxxc5 Ets2 Hoxb3 Pax5 Thra Tcf7l2 Mecom Tal1 Zfp281 Zbtb18 Myc Zfp202 Zfp647* |
| **w6:** $T_{RM}$ | *Nr4a3 Nr4a1 Atf3 Crem Egr1 Klf6 Junb Fosb Tgif1 Isl2 Snai1 Atf4 Irf4 Fosl2 Jun Phox2b Tlx2 Klf5 Cdx1 Mxd1 Shox2 Etv3 Fos Egr2 Nobox Foxi1 Msx1 Vsx1 Wt1 Msx3 Onecut1 Gm28047* |
| **w7:** $T_{CM}$ | *Elk4 Foxo3 Plag1 Zfp341 Usf2 Rel Rreb1 Zbtb33 Pou4f1 Zbtb6 Sp4 Erf Yy1 Foxk1 Zbtb26 Zfp691 Epas1 Cic Rara Arid2 Srebf2 Bach1 Glis3 Zfp398 Jrkl Nfat5 Sox12 Zscan26 Zfp770* |

**e**

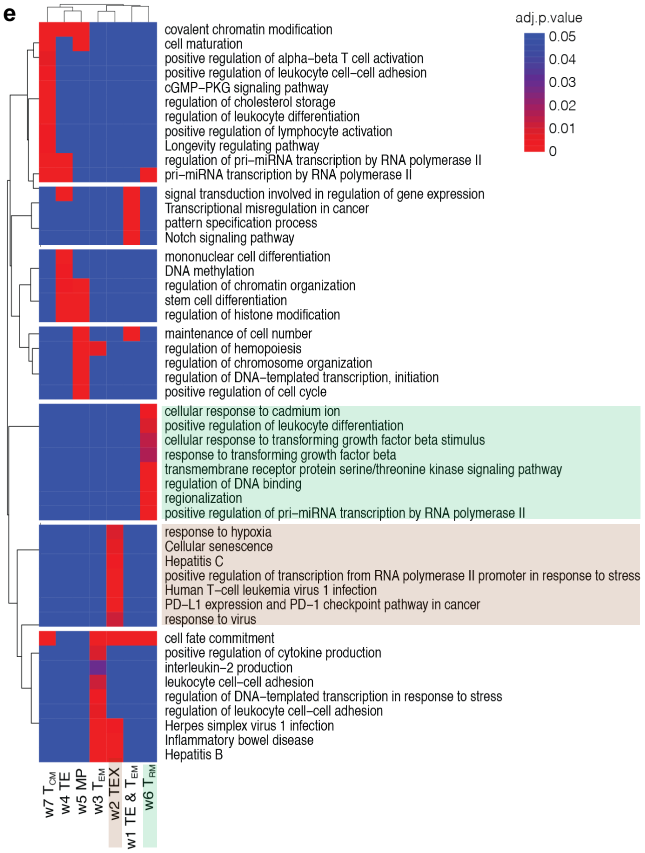


**Extended Data Fig. 3 | TF wave analysis. a,** Schematic of the analysis pipeline. **b,** Selection of algorithms and parameters for TF wave analysis. The Pearson correlation was chosen for the distance metric, with k=7 chosen as the optimal cluster number. **c,** Seven TF waves. Circles represent specific cell states. Red color indicates normalized PageRank scores. **d,** List of TF members in each wave. **e,** Heatmap of biological pathways enriched in each TF wave. Red-blue color scale indicates the p-value.

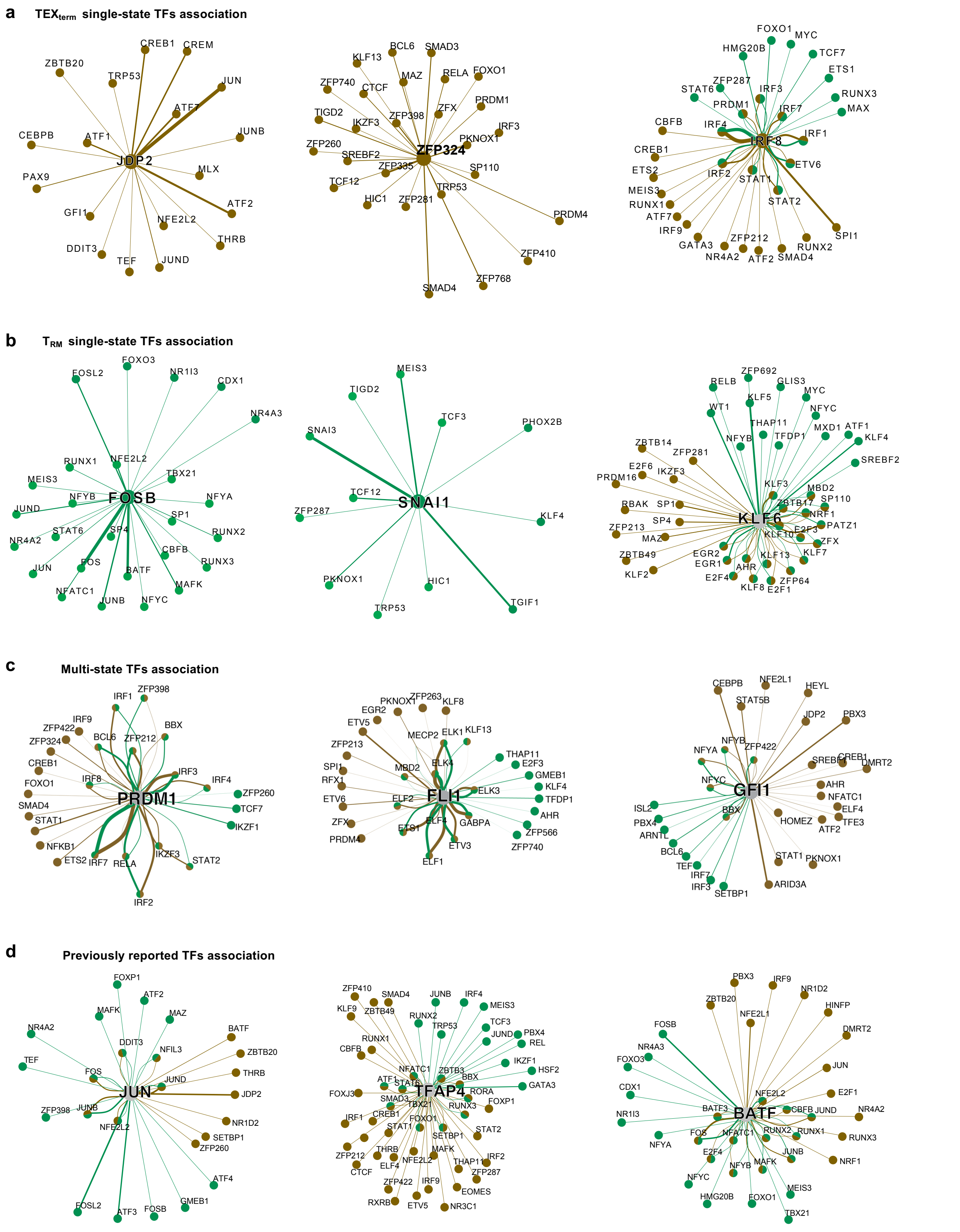


**Extended Data Fig. 4 | TF–TF association network in $TEX_{term}$ and $T_{RM}$ cell states. a–c,** TF–TF associations of (**a**) $TEX_{term}$ single-state TFs (JDP2, ZFP324, and IRF8); (**b**) $T_{RM}$ single-state TFs (FOSB, SNAI1, and KLF6); and (**c**) multi-state TFs shared by $TEX_{term}$ and $T_{RM}$ (PRDM1, FLI1, and GFI1). (**d**) previously reported TFs whose overexpressions prevent $TEX_{term}$–JUN, TFAP4, and BATF. Line color indicates state specificity: $T_{RM}$ (green) or $TEX_{term}$ (brown) state. Line thickness represents interaction intensity.

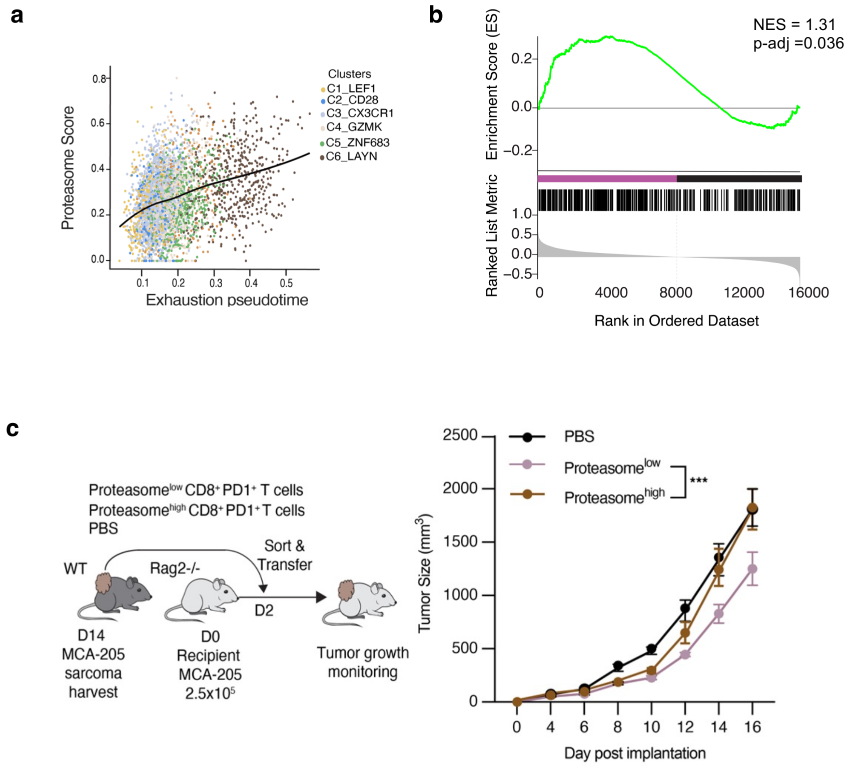


**Extended Data Fig. 5 | TF network analysis reveals proteasome pathway enrichment in $TEX_{term}$ state with diminished tumor control function. a,** Pseudotime analysis of $CD8^+$ TIL scRNA-seq data from NSCLC patients (n = 14), showing a positive correlation between proteasome gene scores (KEGG: M10680) and T cell exhaustion. **b,** Gene set enrichment analysis (GO:0043161, Proteasome-mediated ubiquitin-dependent protein catabolic process) of RNA-seq from $CD8^+$ splenocytes (black) and TILs (purple), n = 3 each. **c,** Tumor growth of $Rag2^{-/-}$ mice bearing MCA-205 sarcomas infused with $proteasome^{high}$ or $proteasome^{low}$ $CD8^+$ TILs isolated from C57BL/6 mice bearing MCA-205 tumors. n = 5 per group. Two-way ANOVA Tukey's multiple comparison test were performed. *****P*<0.0001, ****P*<0.001, ***P*<0.01, **P*<0.05.

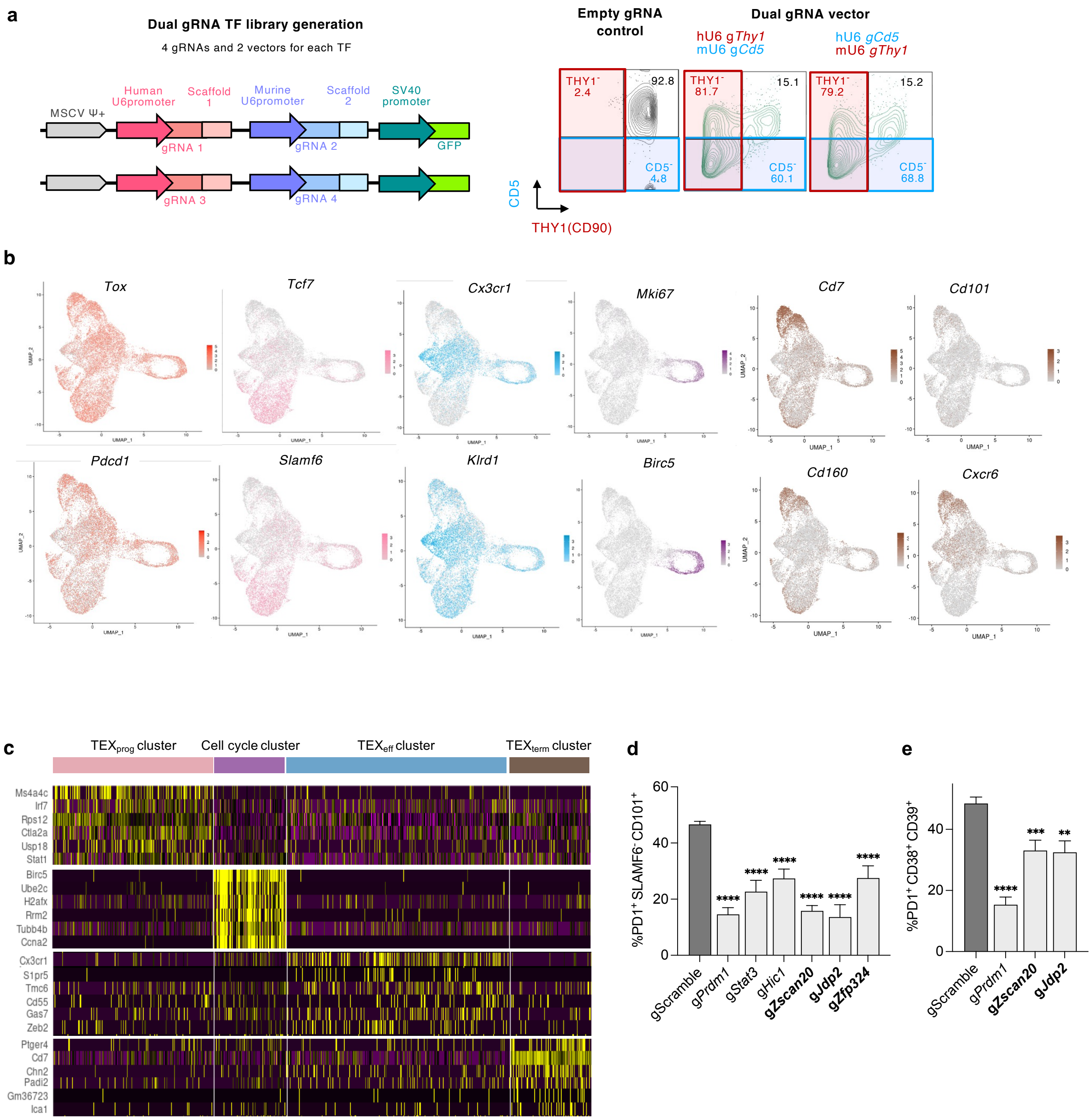


**Extended Data Fig. 6 | *in vivo* Perturb-seq with dual-guide RNA in LCMV chronic infection and individual validation. a,** Retroviral vector design for dual-gRNA delivery. **b,** Feature plot of differentiation markers (Pan-exhaustion: red, $TEX_{prog}$: pink, $TEX_{eff}$: blue, Cell cycle: purple, $TEX_{term}$: brown). **c,** Heatmap of marker gene expression across cell state clusters identified by Seurat's FindMarkers() function. **d,** Frequency of $PD1^+$ $SLAMF6^-$ $CD101^+$ cells (**d**). Frequency of $CD38^+$ $CD39^+$ double-positive cells (**e**). Statistical analysis: Ordinary one-way ANOVA with Dunnett's multiple comparisons test versus gScramble (n ≥ 5 from ≥2 biological replicates). Data are presented as mean ± s.e.m. ****$P < 0.0001$; ***$P < 0.001$; **$P < 0.01$; *$P < 0.05$.

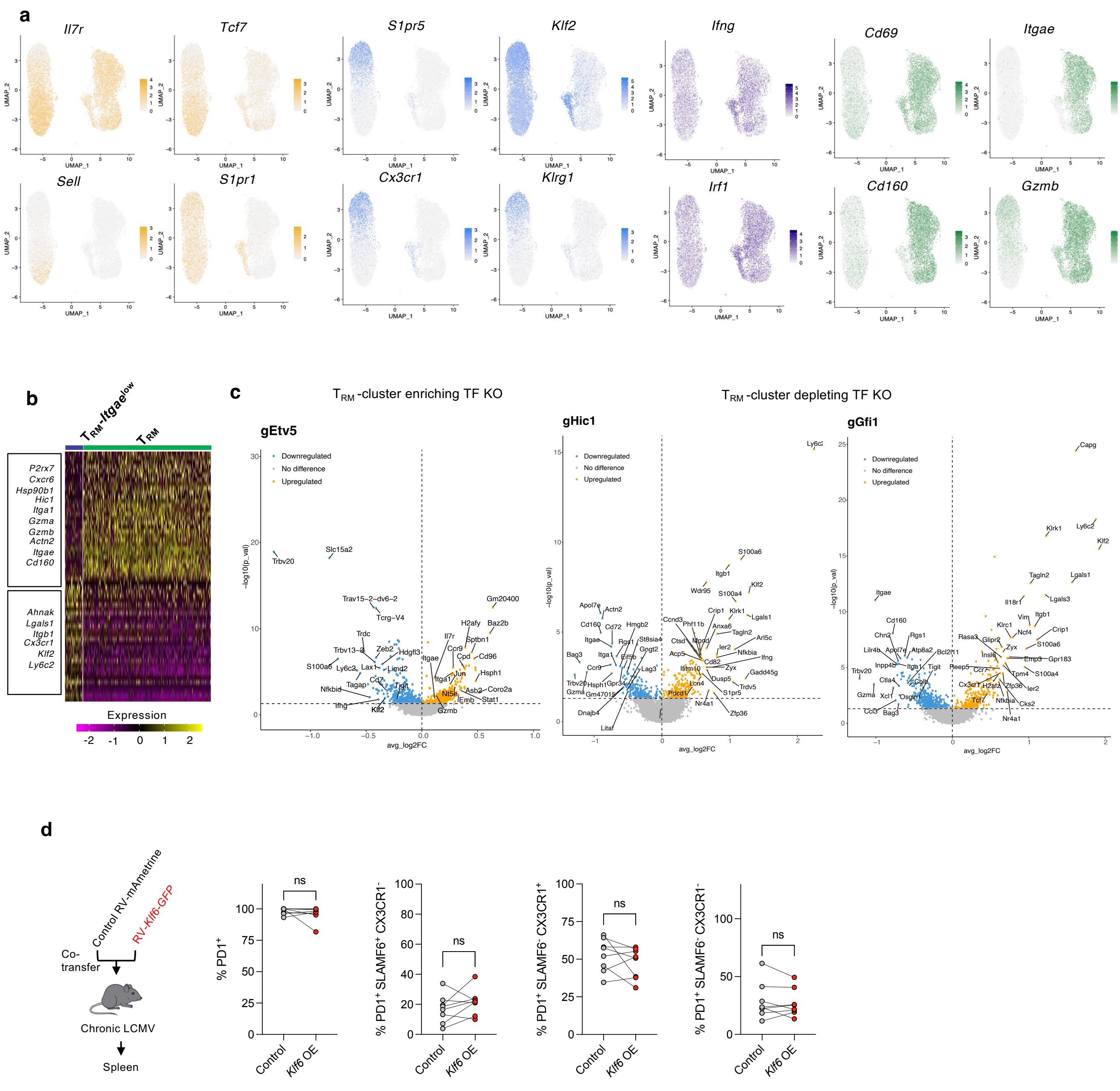


**Extended Data Fig. 7 | Single-cell transcriptomic profiling of CD8$^{+}$ T cells from Perturb-seq in acute LCMV infection.** **a,** Feature plots of differentiation marker genes ($T_{CM}$: yellow, $T_{EM}$: blue, $T_{RM}$-*Itgae*$^{low}$: dark blue, $T_{RM}$: green). **b,** Heatmap of differentially expressed genes between $T_{RM}$-*Itgae*$^{low}$ and $T_{RM}$ clusters. **c,** Volcano plots of differentially expressed genes in Cas9$^{+}$ P14 CD8$^{+}$ T cells expressing *gEtv5*, *gHic1*, or *gGfi1*. **d,** $T_{RM}$ single-state TF, *Klf6* overexpression does not accelerate T cell exhaustion. Experimental setup: *Klf6*-RV or control-RV transduced P14 CD8$^{+}$ T cells co-transferred into mice infected with chronic LCMV-Clone 13. **b-e,** Quantification of the frequency of PD1$^{+}$, $TEX_{prog}$ (PD1$^{+}$ SLAMF6$^{+}$ CX3CR1$^{-}$), $TEX_{eff}$ (PD1$^{+}$ SLAMF6$^{-}$ CX3CR1$^{+}$) and $TEX_{term}$ (PD1$^{+}$ SLAMF6$^{-}$ CX3CR1$^{-}$) populations. Paired t-tests (n ≥ 6 from ≥2 biological replicates). Data are presented as mean ± s.e.m. ****$P$ <0.0001, ***$P$ < 0.001, **$P$ < 0.01, *$P$ < 0.05.

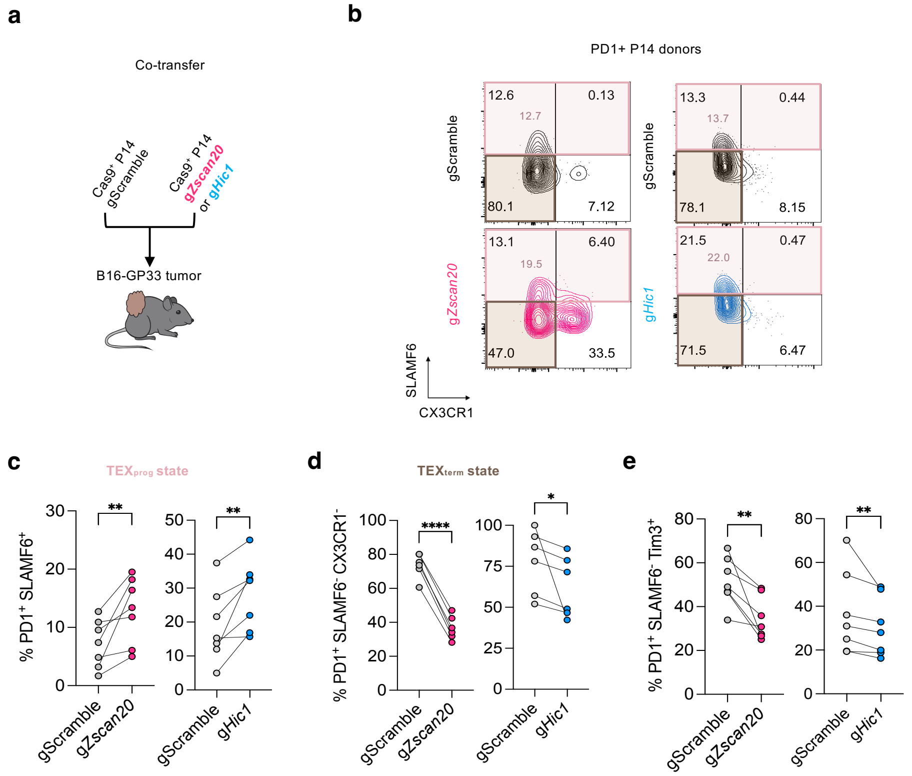


**Extended Data Fig. 8 | Depletion of TEX$_{term}$-single state TF, *Zscan20* and TEX$_{term}$ and T$_{RM}$ multi-state TF, *Hic1* reduces T cell exhaustion. a,** Co-transfer of Cas9$^{+}$ P14 CD8$^{+}$ T cells transduced with RV-g*Zscan20* or RV-g*Hic1*, mixed with gRNA control RV transduced cells into B16-GP33 tumor-bearing mice. **b,** Representative flow plots of PD1$^{+}$ P14 CD8$^{+}$ T cells stained for SLAMF6 and CX3CR1. **c-e,** Quantification of (**c**) TEX$_{prog}$ (PD1$^{+}$ SLAMF6$^{+}$ TIM3$^{-}$), (**d**) TEX$_{term}$ (PD1$^{+}$ SLAMF6$^{-}$ CX3CR1$^{-}$), and (**e**) non-progenitor, exhausted (PD1$^{+}$ SLAMF6$^{-}$ TIM3$^{+}$) popultation. Paired t-tests (n ≥ 6 from ≥2 biological replicates). Data are presented as mean ± s.e.m. ****P < 0.0001, ***P < 0.001, **P < 0.01, *P < 0.05.

**a**

| Author (Year) Accession | Data type | Tumor types | Original Annotated CD8+ T cell clusters included in our meta-analysis |
|---|---|---|---|
| Long (2022) PRJNA768891 | scRNA-seq, scATAC-seq | ccRCC | CD4+ T cell, Treg, CD8+ T cell (further clustered into scRNA-seq: Tissue-resident C2, Tissue-resident C1, Exhaustion, Exhausted IEG; scATAC-seq: Exhausted IEG C2, Exhausted IEG C1, Exhaustion, Tissue-resident, Unknown) |
| Terekhanova (2023) GSE240822 | snRNA-seq, snATAC-seq | ccRCC | CD8+ Naive, CD8+ $T_{CM}$, CD8+ $T_{EM}$, proliferating, MAIT |
| Terekhanova (2023) GSE240822 | snRNA-seq, snATAC-seq | GBM | CD8+ Naive, CD8+ $T_{CM}$, CD8+ $T_{EM}$, proliferating, MAIT |
| Riegel (2023) EGAS00001006141 | scATAC-seq | BCC, HCC, RCC, HNSCC | CD8+ Naive, CD8+ MAIT, CD8+ Heatshock (T_HS), CD8+ Memory, CD8+ Cytotoxic (T_CTX), CD8+ pre-T_Ex, CD8+ prolif-T_Ex, CD8+ term-T_Ex |
| Yost (2019) GSE123814 | scRNA-seq | BCC | Naive T, CD8+ memory, CD8+ activated T, CD8+ $T_{EM}$, exhausted/activated CD8+ , exhausted CD8+ T |
| Ma (2019) GSE125449 | scRNA-seq | HCC | CD8-C1-GZMK, CD8-C2-GZMA, CD8−C3−GNLY |
| Young (2018) Data S1 from PMID:30093597 | scRNA-seq | RCC | CD8+ T cell, Proliferating T cell |
| Cillo (2020) GSE139324 | scRNA-seq | HNSCC | CD8+ T cells |
| Zheng (2021) GSE156728 | scRNA-seq | BC, BCL, CHOL, ESCA, FTC, MM, OV, PACA, RC, THCA, UCEC | CD8.c15 (ISG+ CD8+ T), CD8.c12 (terminal Tex), CD8.c11 (GZMK+ Tex), CD8.c13 (OXPHOS− Tex), CD8.c05 (GZMK+ early Tem), CD8.c10 (ZNF683+CXCR6+ Trm), CD8.c06(GZMK+ Tem), CD8.c14 (TCF7+ Tex), CD8.c17 (NME1+ T), CD8.c07 (Temra), CD8.c01 (Tn), CD8.c09 (KIR+TXK+ NK-like), CD8.c02 (IL7R+ Tm), CD8.c04 (ZNF683+CXCR6− Tm), CD8.c16 (Tc17), CD8.c08 (KIR+EOMES+ NK-like) |

**b** Human Pan-cancer scMultiomics Cluster markers

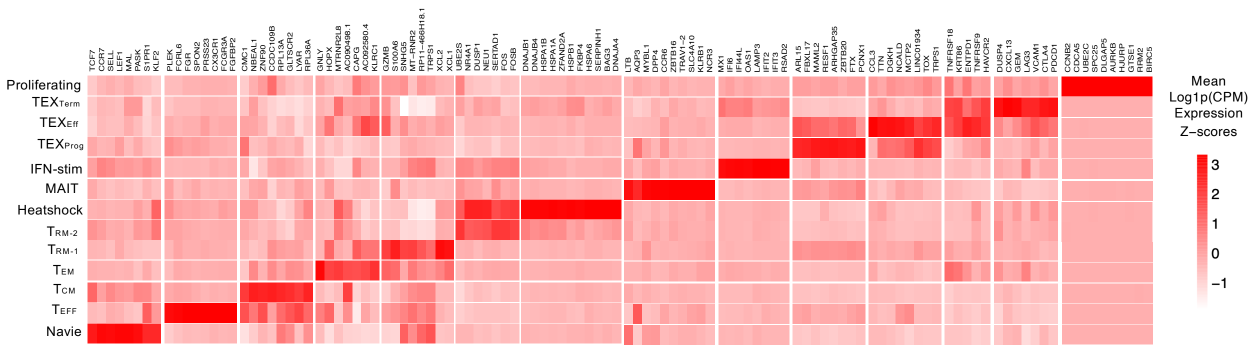


**c** Murine $T_{RM}$ single-state TF activity in human pancancer

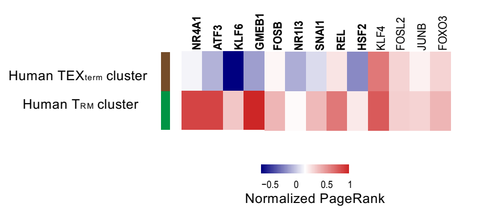


**d** Murine $TEX_{term}$ & $T_{RM}$ multi-state TF activity in human pancancer

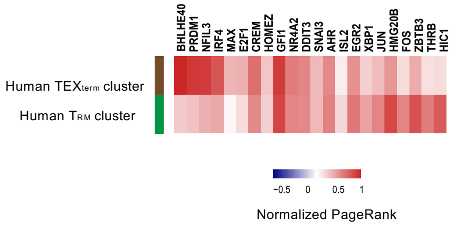


**Extended Data Fig. 9 | Conservation of single- and multi-state TFs in human pan-cancer $TEX_{term}$ and $T_{RM}$ cell states. a,** Human pan-cancer datasets utilized in this study[52–59]. **b,** Cluster-specific marker mRNA expression across integrated pan-cancer single-cell multi-omics datasets. **c,** TF activity analysis using Taiji on matched matched scRNA-seq and scATAC-seq datasets from various human cancers, including ccRCC, GBM, BCC, HNSCC, HCC, and RCC. CD8+ T cell states were annotated using canonical marker gene expression. PageRank scores derived from Taiji were log-transformed, averaged per state, and z-score normalized. Results were visualized with a focus on TF activity $T_{RM}$ and $TEX_{term}$ cell states. **d,** mRNA expression of TFs were compared across human CD8+ T cell states. The dataset encompassed T cells from 15 tumor types. Cell type annotations provided by the original study were retained and mapped to the following broad categories of clusters: $T_{RM}$, $TEX_{term}$, $T_{EM}$, Memory, Naive, and cell cycle. Seurat's AverageExpression function was used, followed by z-score normalization. Data visualization emphasized comparisons between $T_{RM}$ and $TEX_{term}$ cell states. Cell-state-selectivity conserved TFs in humans and mice are highlighted in bold (**c, d**).

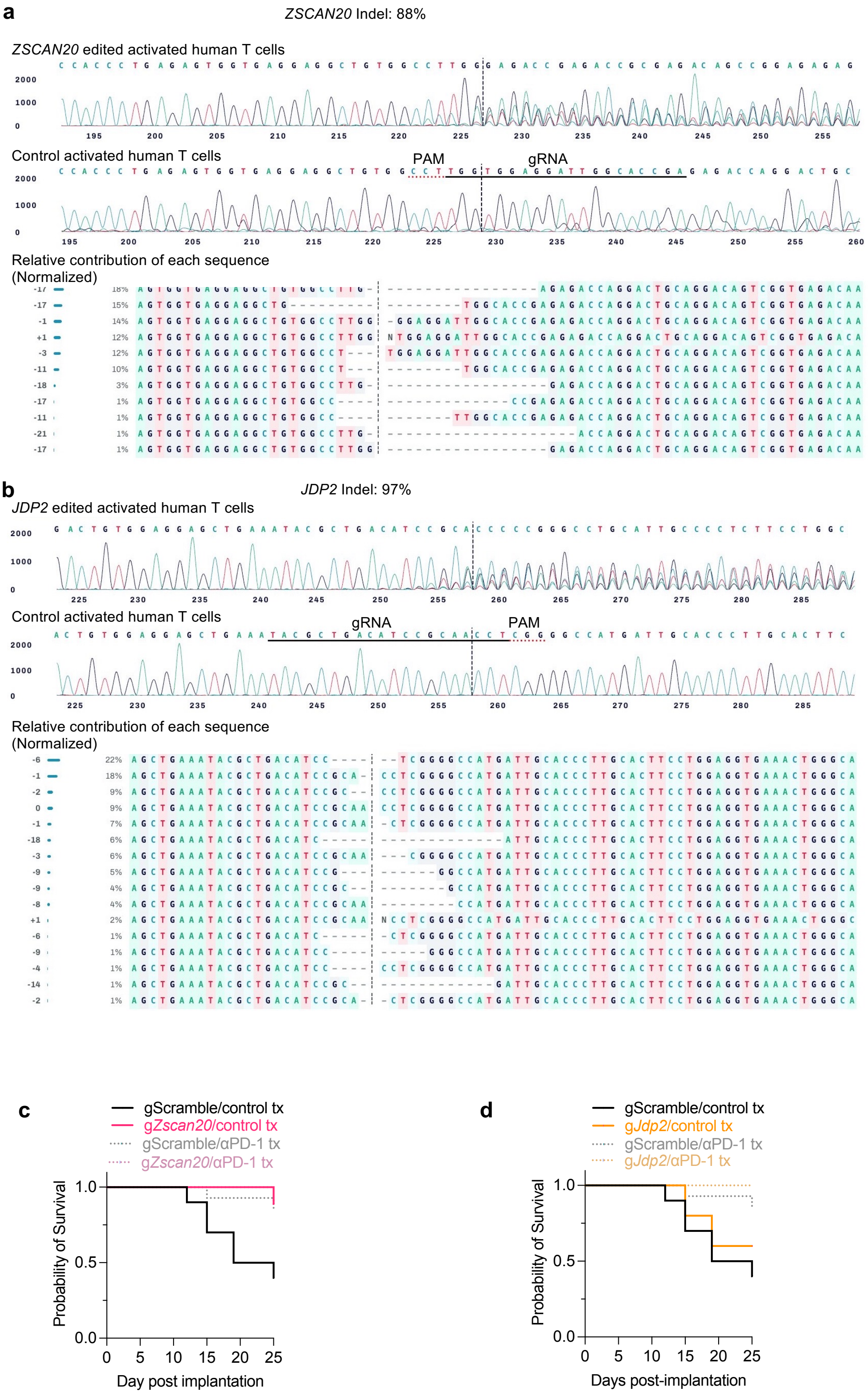


**Extended Data Fig. 10 | CRISPR validation of $TEX_{term}$ single-state TF KO in human PBMCs and antitumor activity of TF-deficient T cells in mice.** Indel frequencies, representative Sanger sequencing traces, and indel distributions for (**a**) *ZSCAN20* and (**b**) *JDP2* KOs. **c,** Survival of B16-GP33 bearing mice receiving *Zscan20* KO or control P14 $CD8^+$ T cells, followed by anti-PD1 or isotype IgG2a treatment. **d,** Survival of tumor-bearing mice with *Jdp2* KO P14 $CD8^+$ T cell transfer.
